# rRNA Biogenesis Regulates Mouse 2C-like State by 3D Structure Reorganization of Peri-Nucleolar Heterochromatin

**DOI:** 10.1101/2021.08.04.454705

**Authors:** Hua Yu, Zhen Sun, Tianyu Tan, Hongru Pan, Jing Zhao, Ling Zhang, Jiayu Chen, Anhua Lei, Yuqing Zhu, Lang Chen, Yuyan Xu, Ming Chen, Shaorong Gao, George Q. Daley, Jin Zhang

## Abstract

Nucleolus is the organelle for ribosome biogenesis and for sensing various types of stress. Its role in regulating stem cell fate is unclear. Here, we present multiple lines of evidence that nucleolar stress induced by interfering rRNA biogenesis can drive 2-cell stage embryo-like (2C-like) transcriptional program and induce an expanded 2C-like cell population in mouse embryonic stem (mES) cells. Mechanistically, the nucleolar integrity mediated by rRNA biogenesis maintains the normal liquid-liquid phase separation (LLPS) of nucleolus and the formation of peri-nucleolar heterochromatin (PNH). Upon rRNA biogenesis defect, the natural LLPS of nucleolus is disrupted, causing dissociation of NCL/TRIM28 complex from PNH and changes of epigenetic states and reorganization of the 3D structure of PNH, which leads to *Dux*, a 2C program transcription factor gene, to be released from the PNH region and activation of 2C-like program. Correspondingly, embryos with rRNA biogenesis defect are incompatible to develop from 2-cell (2C) to 4-cell embryos, with delayed repression of 2C/ERV genes and a transcriptome skewed toward earlier cleavage embryo signatures. Our results highlight that rRNA-mediated nucleolar integrity and 3D structure reshaping of PNH compartment regulates the fate transition of mES cells to 2C-like cells, and that rRNA biogenesis is a critical regulator during the 2-cell-to-4-cell transition of murine pre-implantation embryo development.

## Main

Two-cell (2C) stage embryonic cells are totipotent cells in an earlier stage of embryo development and can generate all cell types of embryonic and extraembryonic tissues. In the culture of mouse embryonic stem (mES) cells, a rare population of mES cells sporadically transit into a 2C stage embryo-like (2C-like) cells with similar molecular features of totipotent 2C-stage embryos ^1–3^. Recent works have demonstrated that the conversion of mES cells to 2C-Like cells is regulated by a variety of factors related to epigenetic modification, including histone methylation and acetylation ^1, 4, 5^ and DNA methylation ^6, 7^. In addition, it was also found that RNA hydroxymethylation ^8^ and protein sumoylation ^9^ can affect the epigenetic state of chromatin to regulate the activation of ERV genes. The structure of chromatin is an important epigenetic factor and is closely related to the regulation of gene expression and cell fate transition. Interestingly, chromatin structure appears to emerge as an important factor in 2C gene regulation and transition of mES cells to 2C-like cells. For instance, 2C/ERV gene activation is regulated by a pioneer transcription factor Dux which increases chromatin accessibility ^10–15^, and two recent studies reported that the pluripotency factors DAPP2 and DAPP4 and the maternal factor NELFA can bind to the promoter region of *Dux* and directly trans-activate its expression ^16, 17^. Moreover, 2C-like cells can be induced by downregulation of chromatin remodeling factor CAF-1 ^18^. However, the molecular players of chromatin structure in mES cell to 2C-like cell transition have yet to be fully understood.

With the emergence and development of high-throughput chromatin conformation capture technology (Hi-C), the dynamic changes of higher-order chromatin structures during early embryonic development and stem cell differentiation have been elucidated ^19–22^. Two-cell embryo or 2C-like cells show contrasting differences from inner cell mass or ES cells ^21, 23^, suggesting that the 3D chromatin structure is a key factor mediating the transition of mES cells to 2C-like cells. Importantly, one of the mechanisms of *Dux* expression is dependent on a complex of nucleolin NCL and heterochromatin factor TRIM28 in the Peri-Nucleolar Heterochromatin (PNH) region^14, 24, 25^. However, it is not completely known how nucleolus integrity influences higher-order chromatin structure and how the chromatin structure determines *Dux* expression.

Here, we found that inhibition of nucleolar rRNA biogenesis triggered nucleolar stress which activated 2C-like transcriptional program and induced an expanded 2C-like cell population in mES cells with a mechanism involving 3D structure reorganization of the PNH and the *Dux* expression. Consistently, 3D structure of PNH reorganizes after early 2-cell during murine early embryo development, which coincides with rRNA biogenesis and Dux repression. Moreover, in mouse early embryos, rRNA biogenesis and matured nucleolus are indispensable for the 2-cell to 4-cell transition. Taken together, our findings for the first time provided a novel mechanistic perspective of rRNA biogenesis in regulating the homeostasis between 2C-like and mES cells and highlighted that rRNA biogenesis in the nucleolus is a critical molecular switch from ZGA gene expressing 2-cell stage to nucleolus-matured blastocyst stage embryos.

## Results

### Inhibition of rRNA biogenesis activated the 2C-like transcriptional program and induced an expanded 2C-like cell population in mES cells

We first explored whether nucleolar stress produced by inhibiting rRNA biogenesis could induce cell fate reprogramming to 2C-like cells (2CLCs) by performing RNA-seq analysis of mES cells treated by three inducers of cellular stress, including CX-5461, an RNA polymerase I (Pol I) inhibitor; rotenone, an electron transport chain complex 1 inhibitor, and rapamycin, a mTOR pathway inhibitor (CX-5461 treatment dosage: 2uM, CX-5461 treatment time: 12h; rotenone treatment dosage: 1uM, rotenone treatment time: 12h; rapamycin treatment dosage: 2uM, rapamycin treatment time: 12h). We found that nucleolar stress induced by CX-5461^26, 27^ activated the 2-cell marker genes *Zscan4d*, *Dux* and *Gm12794* and repressed pluripotent marker gene *Pou5f1*. However, the other two cellular stresses upon rotenone or rapamycin treatment did not influence the expression of these genes (Fig.1a-1b and Extended Data Fig.S1a-S1b). The 2C-like transcriptional program was characterized by activation of transposable elements (TEs), particularly major satellite repeats (GSAT_MM) and ERVL subclasses MERVL-int and MT2_Mm. We systematically examined ERV genes and found global up-regulation of GSAT_MM and each LTR class (Fig.1C and Extended Data Fig.S1e), particularly GSAT_MM, MERVL-int and MT2_Mm sub-classes, in CX-5461-treated mES cells (Fig.1c and Extended Data Fig.S1c-S1d). However, the other two types of stress, did not activate these repeat elements (Fig.1c and Extended Data Fig.S1c-S1d). Using unsupervised K-means clustering analysis, we identified four gene clusters specifically expressed in different stages during mouse pre-implantation embryo development (Extended Data Fig.S1f)^28^. We found that CX-5461 treatment upregulated 2-cell expressing cluster 1 (C1) and 2-cell/4-cell expressing cluster 2 (C2) genes, and decreased genes expressed in other two stages (C3 and C4) (Fig.1d and Extended Data Fig.S1f). Yet, other two cellular stresses did not induce this expression pattern (Fig.1d and Extended Data Fig.S1f). Moreover, unsupervised hierarchical clustering of transcriptomes of pre-implantation embryos and mES cells from published studies of 2C-like cells confirmed that mES cells treated with CX-5461 were most like the sorted 2CLCs from mES cells, or genetically modified mES cells with 2CLC signatures from other studies as well as the 2C embryos^1, 11, 14, 16–18, 29–31^ (Fig.1e). As expected, we observed the abundance of rRNA is significantly reduced under CX-5461 treatment (Extended Data Fig.S1g). Together, these results demonstrate that nucleolar stress induced by rRNA biogenesis defect activated 2C-like transcriptional program in mES cells.

**Fig.1:**
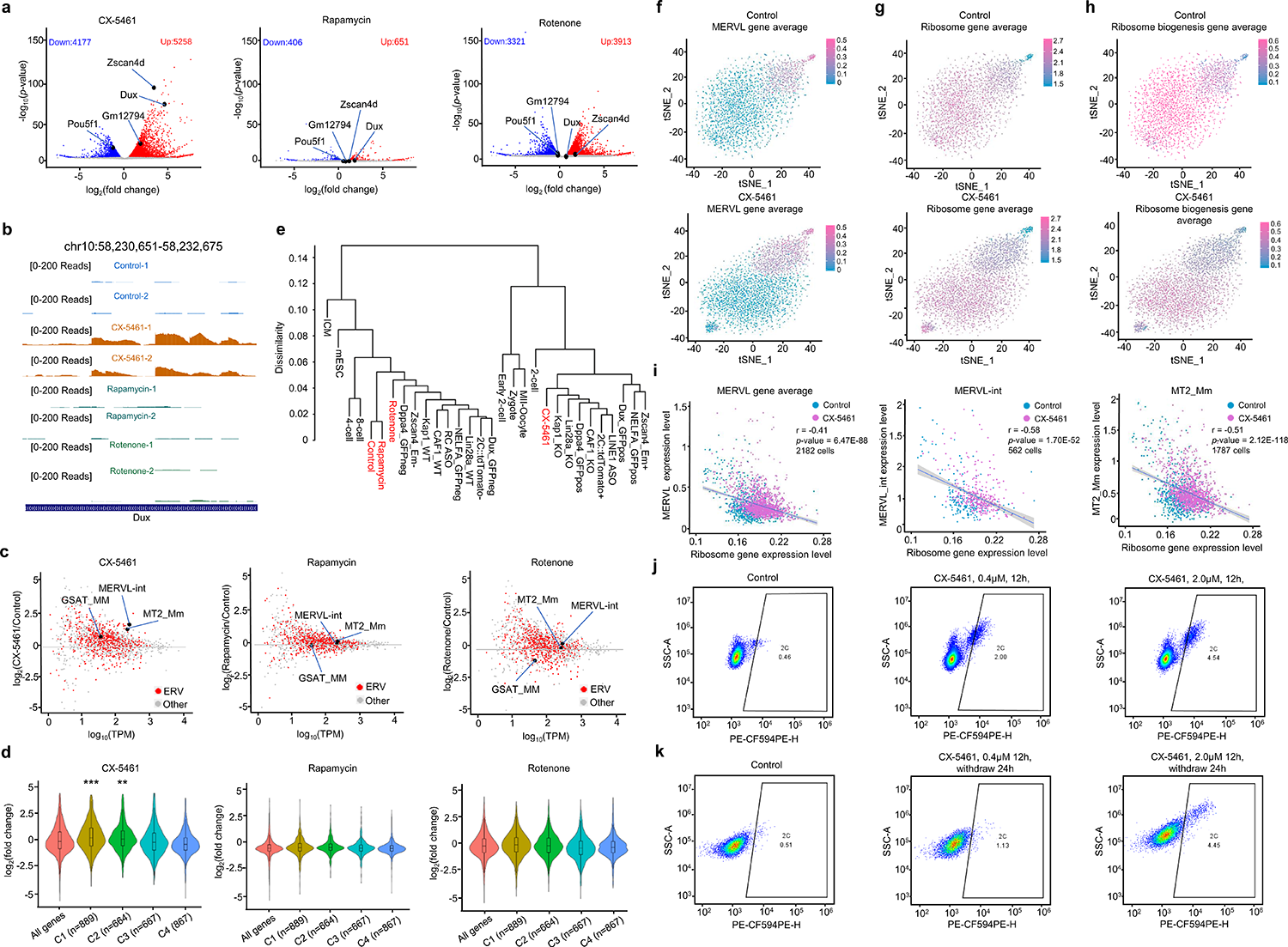
Inhibition of rRNA biogenesis activated 2C-like transcriptional program and induced an expanded 2C-like cell population in mES cells. **a)** Volcano plots of RNA-sequencing data comparing gene expression of control and cellular stress inducer-treated mES cells (GEO accession GSE166041). **b)** UCSC Genome Browser viewing of RNA-sequencing results at the *Dux* locus (GEO accession GSE166041). **c)** MA plots of RNA-sequencing data comparing repeat sequence expression of control and cellular stress inducer-treated mES cells (GEO accession GSE166041). **d)** Violin plots demonstrating the expressional changes of stage-specific gene clusters of mouse pre-implantation embryos under different types of cellular stress treatment; **: p<0.01, ***: p<0.001, Mann-Whitney U test (GEO accession GSE166041). **e)** Hierarchical clustering of transcriptomes from our study, published 2C-like cell model studies and pre-implantation mouse embryos; Control, CX-5461, Rotenone and Rapamycin (GEO accession GSE166041), 2C::tdTomato+ and 2C::tdTomato-(GEO accession GSE33923); Zscan4_Em+ and Zscan4_Em-(GEO accession GSE51682); Kap1_KO and Kap1_WT (GEO accession GSE74278); CAF1_WT and CAF1_KO (GEO accession GSE85632), Dux_GFPpos and Dux_GFPneg (GEO accession GSE85632); LINE1 ASO and RC ASO (GEO accession GSE100939); Dppa4_GFPpos and Dppa4_GFPneg (GEO accession GSE120953), NELFA_GFPpos and NELFA_GFPneg (GEO accession GSE113671); Lin28a_KO and Lin28a_WT (GEO accession GSE164420); MII-Oocyte, Zygote, Early 2-cell, 2-cell, 4-cell, 8-cell, ICM and mES cells (GEO accession GSE66390). **f)** tSNE feature plots demonstrating the averaged expression levels of *MERVL* genes in 2981 control mES cells and 3219 CX-5461-treated mES cells. **g)** tSNE feature plots demonstrating the averaged expression levels of ribosome genes in 2981 control mES cells and 3219 CX-5461-treated mES cells. **h)** tSNE feature plots demonstrating the averaged expression levels of ribosome biogenesis genes in 2981 control mES cells and 3219 CX-5461-treated mES cells (GEO accession GSE166041). **i)** Scatter plots demonstrating negative correlation of expression level between *MERVL/MERVL-int/MT2_Mm* and ribosome genes; Each dot represents a single cell with detectable ERV expression; *r* denotes correlation coefficient; *p*-value was obtained by cor.test function in R software (GEO accession GSE166041). **j)** FACS analysis on 2C::tdTomato+ mES cells upon different treatment doses of CX-5461, showing the change of percentage of 2C-like cells. **k)** FACS analysis on 2C::tdTomato+ mES cells after 12 hour treatment and 24 hour withdrawal of CX-5461, showing the change of percentage of 2C-like cells.

We next asked how the population homeostasis of 2CLCs and ES cells altered in response to rRNA biogenesis defect at the single cell level. In line with bulk RNA-seq data, we observed that the 2C marker genes are up-regulated and pluripotent genes were downregulated in CX-5461-treated mES cells (Extended Data Fig.S1h-S1j). As expected, we found a marked expansion of the population with MERVL expression in mES cells treated with CX-5461 (Fig.1f). Strikingly, we observed that the expression level of MERVL genes showed significant negative correlation with ribosomal protein genes (Fig.1g-1i and Extended Data Fig.S1k). Using a 2C::*tdTomato* reporter in which a *tdTomato* gene is under control of a MERVL promoter, we examined 2C status of individual cells by Fluorescence Activated Cell Sorting (FACS) analysis. Consistent with single-cell RNA-seq data, we observed that CX-5461-treatment induced a significant increase in the number of tdTomato positive (2C::tdTomato+) mES cells in a dose-dependent manner (Fig.1j and Extended Data Fig.S1l). Moreover, the percentage of tdTomato positive cells was largely maintained even at 24 hours after CX-5461 withdrawal (Fig.1k). Importantly, although CX-5461, mostly at a high concentration, induced mild cell apoptosis, the majority of tdTomato positive cells were negative for the apoptosis markers Annexin-V and DAPI^32^ (Extended Data Fig.S1m-S1o), suggesting that the emergence of 2C::tdTomato+ cells under CX-5461 treatment was not due to the activation of apoptosis pathway. Collectively, these results demonstrate that inhibiting rRNA biogenesis induced a shift of the ES cell homeostasis toward the MERVL-expressing and ribosomal gene repressed 2CLCs ^33, 34^.

### Deficiency of rRNA biogenesis disrupted normal nucleolar LLPS and epigenetic state of PNH

As it has been reported that ribosomal RNA plays a critical role in maintaining phase separation of nucleolus^35–37^ and phase-to-phase transition is involved in nucleolar stress^37, 38^, we examined whether nucleolar stress induced by rRNA biogenesis defect leads to changes of nucleolar phase separation. Using electron microscopy, we observed that CX-5461-treated mES cells displayed abnormal nucleolar structure, missing the outer layer usually associated with dense electron intensity (Fig.2a). Immunofluorescence of key granular compartment (GC) and nucleolar dense fibrillar component (DFC) marker proteins NPM1 and NCL revealed that CX-5461 treatment led to disappearance of the NPM1- and NCL-marked “ring” structure (Fig.2b-2c). Immunofluorescence of FBL and RPA194 protein also showed abnormal localization with aggregated pattern in the nucleolus upon treatment (Fig.2d-2e). Consistently, Fluorescence Recovery After Photobleaching (FRAP) analysis revealed that CX-5461 treatment markedly increased the mobility of NCL, NPM1 and FBL (Fig.2f-2h). Together, these data demonstrate that nucleolar stress caused disrupted assembly of the phase-separated nucleolar sub-compartments, which became fused and more dynamic liquid-like droplets^37, 38^.

**Fig.2:**
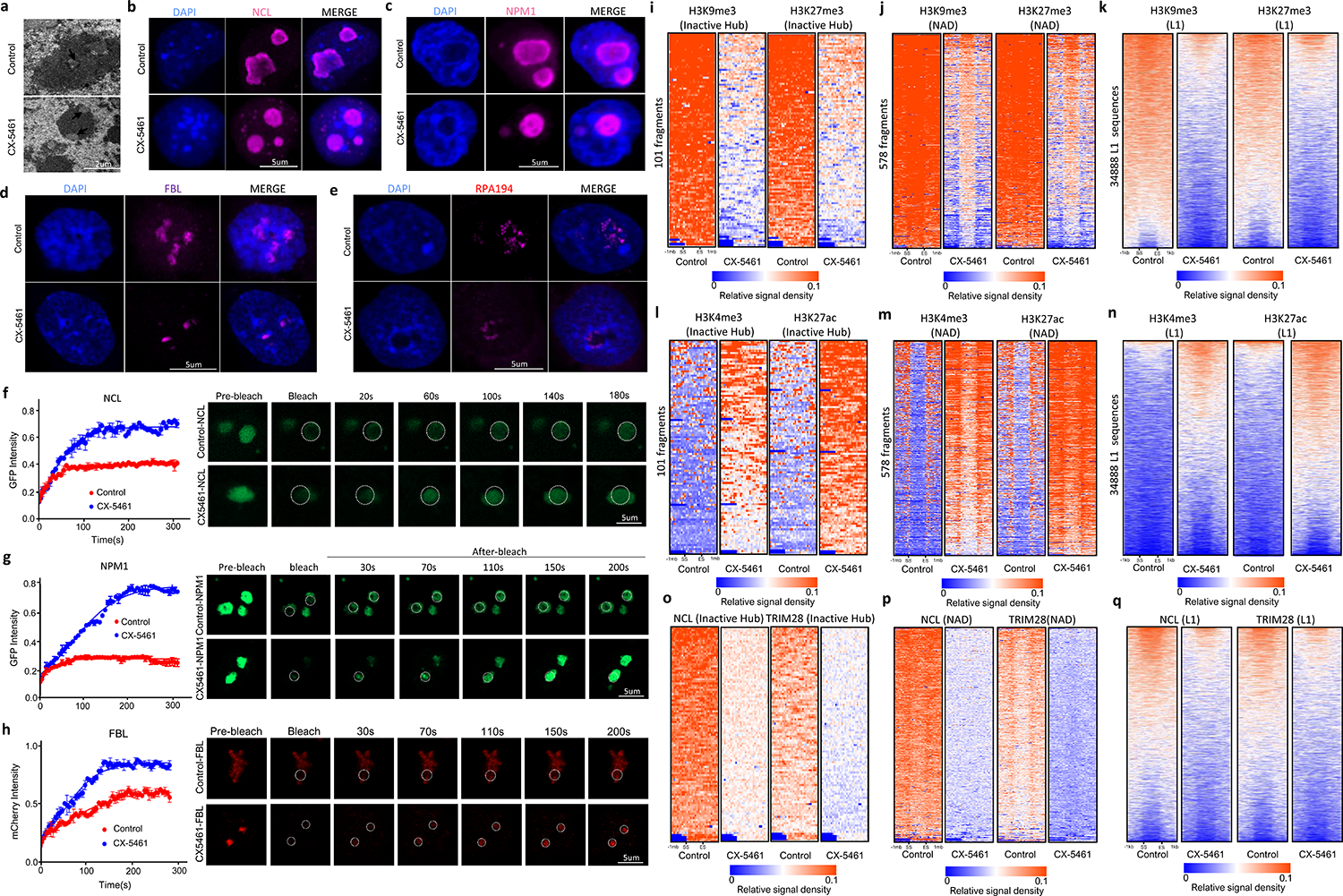
Deficiency of rRNA biogenesis disrupted normal nucleolar LLPS and epigenetic state of PNH region. **a)** CX-5461 treatment causes abnormal nucleolar structure with electron microscopy. **b)** Immunofluorescence staining of NCL in control mES cells and CX-5461-treated mES cells. **c)** Immunofluorescence staining of NPM1 in control mES cells and CX-5461-treated mES cells. **d)** Immunofluorescence staining of FBL in control mES cells and CX-5461 treated mES cells. **e)** Immunofluorescence staining of RPA194 in control and CX-5461 treated mES cells. **f)** FRAP analysis showing CX-5461 treatment causes accelerated recovery after photobleaching of NCL. Shown images are representative of 4 times of experiments. **g)** FRAP analysis showing CX-5461 treatment causes accelerated recovery after photobleaching of NPM1. Shown images are representative of 4 times of experiments. **h)** FRAP analysis showing CX-5461 treatment causes accelerated recovery after photobleaching of FBL. Shown images are representative of 4 times of experiments. **i)** Heatmap plots demonstrate the levels of H3K9me3 and H3K27me3 on within 1mb region around start and end sites of Inactive Hub. The regions of different lengths of Inactive Hub fragments were fitted to 1mb (GEO accession GSE166041). **j)** Heatmap plots demonstrate the levels of H3K9me3 and H3K27me3 within 1mb region around start and end sites of NAD. The regions of different lengths of NAD fragments were fitted to 1mb (GEO accession GSE166041). **k)** Heatmap plots demonstrate the levels of H3K9me3 and H3K27me3 within 1kb region around start and end sites of L1 (GEO accession GSE166041). The regions of different lengths of L1 sequences were fitted to 1kb. **l)** Heatmap plots demonstrate the level of H3K4me3 and H3K27ac within 1mb region around start and end sites of Inactive Hub (GEO accession GSE166041). The regions of different lengths of Inactive Hub fragments were fitted to 1mb. **m)** Heatmap plots demonstrate the levels of H3K4me3 and H3K27ac within 1mb region around start and end sites of NAD. The regions of different lengths of NAD fragments were fitted to 1mb (GEO accession GSE166041). **n)** Heatmap plots demonstrate the level of H3K4me3 and H3K27ac on within 1kb region around start and end sites of L1 (GEO accession GSE166041). The regions of different lengths of L1 sequences were fitted to 1kb. **o)** Heatmap plots demonstrate the level of NCL and TRIM28 within 1mb region around start and end sites of Inactive Hub (GEO accession GSE166041). The regions of different lengths of Inactive Hub fragments were fitted to 1mb. **p)** Heatmap plots demonstrate the levels of NCL and TRIM28 within 1mb region around start and end sites of NAD. The regions of different lengths of NAD fragments were fitted to 1mb (GEO accession GSE166041). **q)** Heatmap plots demonstrate the level of NCL and TRIM28 within 1kb region around start and end sites of L1 (GEO accession GSE166041). The regions of different lengths of L1 sequences were fitted to 1kb.

As it has been observed that phase separation can regulate the epigenetic state of chromatin^39–41^, we examined the epigenetic changes at the loci of the peri-nucleolar heterochromatin (PNH) region in CX-5461-treated mES cells. It has been reported that transcriptionally inactive genomic regions organize into Inactive Hubs around the nucleolus^42^. In addition, the Nucleolar Associated Domains (NAD) or LINE1/L1 repeat sequence regions are also defined as repressive chromosomal segments enriched with peri-nucleolar heterochromatin^43–49^. We thus examined the epigenetic changes on these regions and found decreased H3K9me3 and H3K27me3 levels in CX-5461 treated cells (Fig.2i-2k, Extended Data Fig.S2a and Extended Data Fig.S2b). Moreover, we observed increased H3K4me3 and H3K27ac levels and improved chromatin accessibility at Inactive Hub^42^, NAD^43, 45^ and L1 regions (downloaded from UCSC Table Browser) in CX-5461 treated cells (Fig.2l-2n, Extended Data Fig.S2c-S2g). Previous work has reported that nucleolar protein nucleolin NCL and its interacting partner the heterochromatin protein TRIM28 repress 2C-like program by maintaining the PNH region in mES cells^14^. The disappeared NCL-marked “ring” structure suggested that rRNA biogenesis defect promoted the dissociation of NCL/TRIM28 complex from PNH region. To validate this, we conducted ChIP-seq experiment to investigate the binding changes of NCL/TRIM28 complex on the loci of the PNH region in CX-5461-treated mES cells and found decreased binding of NCL and TRIM28 proteins on Inactive Hub, NAD and L1 regions (Fig.2o-2q, Extended Data Fig.S2e and Extended Data Fig.S2f). Altogether, these results demonstrated the rRNA biogenesis defect affected the normal nucleolar phase separation and changed the epigenetic state of the heterochromatic regions at the periphery of nucleolus by breaking up the binding of NCL/TRIM28 complex on the PNH region.

### 2C/ERV genes were activated through Dux

Recent studies have reported that a pioneer transcription factor, the DUX protein, directly binds to promoters and LTR elements on 2C genes and repetitive elements and activates their transcription^10–12^. As *Dux* expression has been reported to be influenced by nucleolar protein NCL^14^, we speculated that *Dux* is the key molecular regulator for nucleolar stress-mediated activation of 2C-like transcriptional program. To this end, we performed the binding motif sequence enrichment analysis of transcription factors on 5258 genes induced by CX-5461. We found that the significantly enriched motifs include both p53 binding sites and Dux binding sites (Fig.3a). In line with this, we found that 1229 CX-5461-induced genes are p53 direct target genes (Fig.3b, hypergeometric test, p-value=0)^50^, consistent with the fact that p53 signaling is usually activated under nucleolar stress^51–57^. We further analyzed the overlap between CX-5461 treatment-induced genes and *Dux* over-expression-induced genes using published RNA-seq data ^11^, and found 621 genes are overlapped (Fig.3b, hypergeometric test, p-value= 2.96e-195). Using p53 and Dux ChIP-seq data in mES cells ^11, 50^, we further observed that both p53 and Dux showed a pattern of binding to the transcriptional start site (TSS) of CX-5461-induced genes (Fig.3c). The binding pattern of Dux on these genes is weaker when comparing with the strong binding of p53 on CX-5461-induced genes. This was expected as CX-5461 induces many genes that are p53 target, but not Dux target. Interestingly, we found that p53 favors to bind specifically to CX-5461-induced genes, while Dux exclusively binds to the commonly induced genes between CX-5461-treated and *Dux*-overexpressed cells (Fig.3d), and to ERV genes induced by CX-5461 (Fig.3e-3f). Consistently, a significant increase of chromatin accessibility in the promoter region is not observed for 4637 CX-5461-induced genes but is observed for 621 commonly induced genes or 10173 CX-5461-induced ERV genes in *Dux*-overexpressed mES cells ^11^ (Extended Data Fig.S3a). In addition, we observed the decreased H3K9me3 and H3K27me3 levels and increased H3K4me3 and K3K27ac levels around Dux locus (Extended Data Fig.S3b). These analyses suggested that nucleolar stress-induced 2C activation is through *Dux*. To validate this hypothesis, we silenced *Dux* expression in CX-5461-treated mES cells and found that it reversed the 2C/ERV gene induction (Fig.3g). We further performed ChIP-qPCR experiments to assess the changes of H3K9me3 & H3K27me3 levels and Dux binding of CX-5461 induced 2C marker genes after Dux silencing. We observed that, after Dux silencing, both H3K9me3 and H3K27me3 are increased, in contrast, Dux binding is decreased for CX-5461 induced 2C genes (Fig 3h-3j). When the Dux silenced mES cells are treated with CX-5461, we observed that H3K9me3 and H3K27me3 levels, and Dux binding on CX-5461 induced 2C genes are reversed (Fig 3h-3j). We also performed ATAC-seq experiment to assess the changes of chromatin accessibility after Dux silencing. In well support with these results, we observed that chromatin accessibility of CX-5461 induced 2C genes is decreased in Dux silenced mES cells, and this reduction is reversed by CX-5461 treatment (Fig 3k). These results demonstrated that nucleolar stress induced 2C/ERV gene activation through a Dux. In line with Dux’s role in chromatin accessibility, we observed that the 621 commonly induced genes and 10173 CX-5461 induced ERV genes have increased chromatin accessibility, and decreased H3K9me3 and H3K27me3 levels when comparing CX-5461 treated mES cells with control mES cells or comparing 2-cell embryos with ICM stage embryos ^28, 29, 58^ (Extended Data Fig.S3c-S3h). In addition, we also found increased H3K4me3 and H3K27ac levels of 621 commonly induced genes and 10173 CX-5461 induced ERV genes in CX-5461 treated mES cells (Extended Data Fig.S3i-S3j). Collectively, these results demonstrated that nucleolar stress induced a 2C-like transcriptional and epigenetic program in mES cells through Dux.

**Fig.3:**
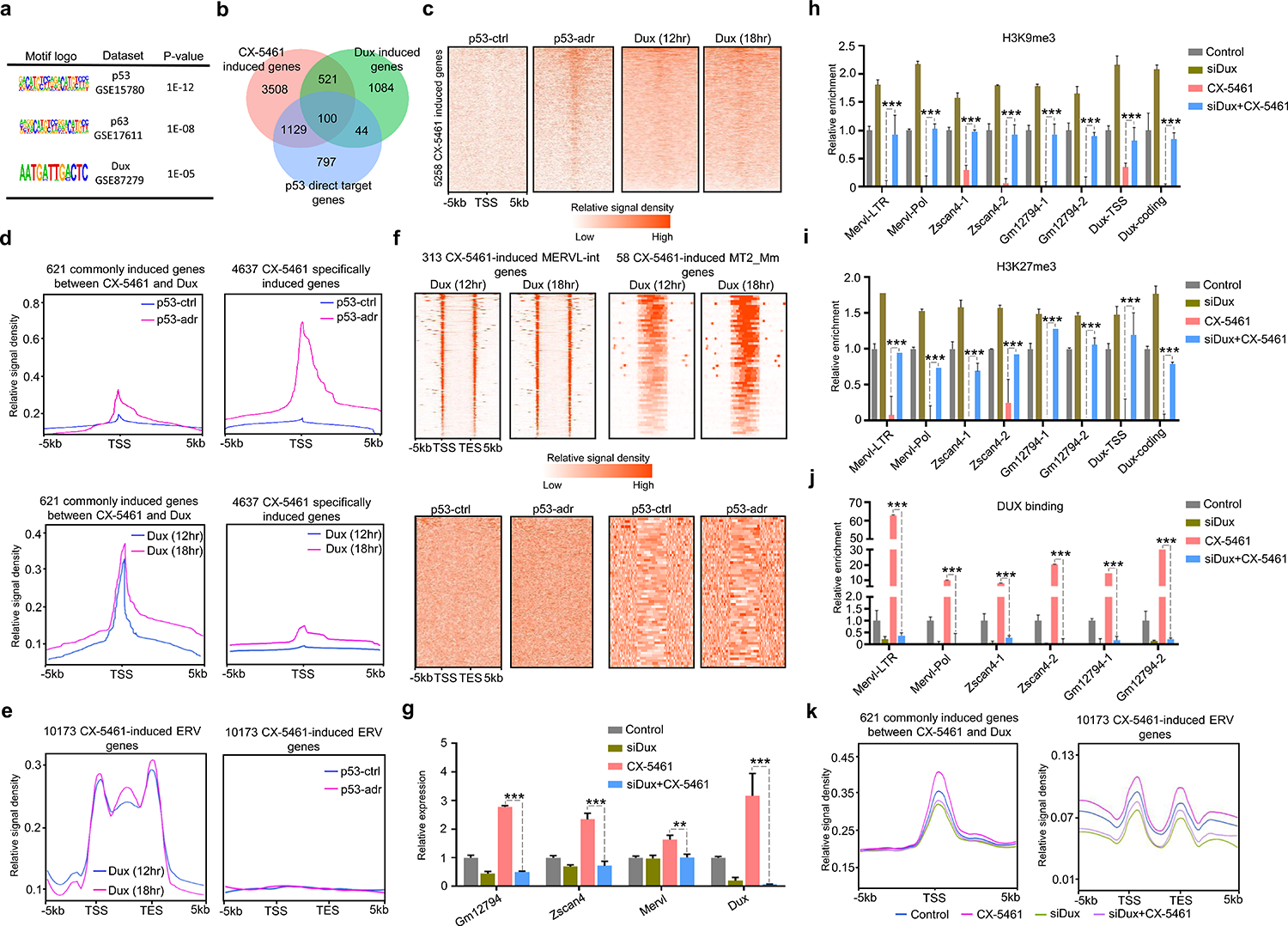
2C/ERV genes were activated through the Dux. **a)** Enriched binding motifs of 5258 genes induced by CX-5461 treatment. **b)** Venn diagrams showing the overlap among CX-5461 treatment induced genes, p53 activated direct target genes and *Dux*-overexpression induced genes. **c)** Heatmap plots demonstrate the binding of p53 and Dux proteins on within 5kb region around transcription start sites of 5258 CX-5461 induced genes using published p53 ChIP-seq data and Dux ChIP-seq data (GEO accession GSE26360 for p53 and GEO accession GSE85632 for Dux). **d)** Line plots demonstrate the meta-analysis results of p53 and Dux protein on within 5kb region around transcription start sites of 621 commonly induced genes between CX-5461 treatment and *Dux* overexpression and 4637 specifically induced genes by CX-5461 using published p53 ChIP-seq data and Dux ChIP-seq data (GEO accession GSE26360 for p53 and GEO accession GSE85632 for Dux). **e)** Line plots demonstrate the meta-analysis results of p53 and Dux protein on within 5kb region around transcription start and end sites of 10173 CX-5461 induced ERV genes using published p53 ChIP-seq data and Dux ChIP-seq data. The regions of different lengths of gene body were fitted to 5kb (GEO accession GSE26360 for p53 and GEO accession GSE85632 for Dux). **f)** Heatmap plots demonstrate the binding of p53 and Dux proteins on within 5kb region around transcription start and end sites of 5258 CX-5461 induced *MERVL-int* and *MT2_Mm* genes; The regions of different lengths of gene body were fitted to 5kb (GEO accession GSE26360 for p53 and GEO accession GSE85632 for Dux). **g)** qRT-PCR showing the expression of *Dux* or 2C-related genes in *Dux* silenced mES cells; **: p<0.01, ***: p<0.001, two-way ANOVA, the replicates of experiment n=3; error bar: standard error of the mean. **h)** ChIP-PCR showing H3K9me3 levels of Dux or 2C-related genes in Dux silenced mES cells; *: p<0.05, **: p<0.01, ***: p<0.001, two-way ANOVA, the replicates of experiment n=3, error bar: standard error of mean. **i)** ChIP-PCR showing H3K27me3 levels of Dux or 2C-related genes in Dux silenced mES cells; *: p<0.05, **: p<0.01, ***: p<0.001, two-way ANOVA, the replicates of experiment n=3, error bar: standard error of mean. **j)** ChIP-PCR showing DUX protein binding levels on 2C-related genes in Dux silenced mES cells; *: p<0.05, **: p<0.01, ***: p<0.001, two-way ANOVA, the replicates of experiment n=3, error bar: standard error of mean. **k)** Line plots demonstrate the meta-analysis results of chromatin accessibility in Dux silenced mES cells within 5kb region around transcription start sites or transcription start and end sites of 621 commonly induced genes between CX-5461 treatment and *Dux* overexpression and 10173 CX-5461 induced ERV genes using published ATAC-seq data. The regions of different lengths of ERV genes were fitted to 5kb (GEO accession GSE166041). p53-ctrl: untreated mES cells, p53-adr: mES cells treated with adriamycin, a DNA damage agent widely used to activate p53, Dux (12h): mES induced with doxycycline for 12 hours, Dux (18h): mES induced with doxycycline for 18 hours.

### rRNA biogenesis defect drove 3D chromatin structure reorganization of PNH and MERVL regions towards the 2C-like state

The disassembly of PNH region suggested that the 3D chromatin structure within PNH region might have reshaped under CX-5461 treatment. To explore the reorganization of 3D chromatin conformation landscape of CX-5461-treated mES cells relative to control mES cells, we performed *in situ* Hi-C with more than four hundred million sequenced raw read pairs per sample. We observed obviously decreased higher-order chromatin interactions within PNH region indicated by the Inactive Hub, NAD and L1 regions in the treated cells (Fig.4a-4c, compared with randomly selected genomic regions, Mann-Whitney U test, the replicates of experiment n=10, averaged *p*-values=0, 0, 3.42E-06 for Inactive Hub, NAD and L1, respectively). Moreover, the *Dux* locus is significantly further away from the PNH region as characterized by the largely decreased Hi-C contacts (Fig.4a-4c and Fig.4g-4i, compared with randomly selected genomic regions, Mann-Whitney U test, the replicates of experiment n=10, averaged *p*-values=1.65E-05, 7.82E-05, 2.68E-03 for Inactive Hub, NAD and L1, respectively). We further analyzed the 3D chromatin structural correlation within PNH region, and between the *Dux* locus and PNH region by comparing Hi-C pearson correlation coefficient (PCC) matrix of control and CX-5461-treated mES cells. We observed the obviously decreased 3D chromatin structural correlation within PNH region (compared with randomly selected genomic regions, Mann-Whitney U test, the replicates of experiment n=10, averaged *p*-values=0 for all Inactive Hub, NAD and L1) and between the *Dux* locus and PNH region (compared with randomly selected genomic regions, Mann-Whitney U test, the replicates of experiment n=10, *p*-values=1.86E-6, 5.03E-6, 4.02E-5 for Inactive Hub, NAD and L1, respectively) (Fig.4d-4f and Fig.4j-4l). We further validated the above findings using DNA Fluorescence in Situ Hybridization (FISH) and found that the *Dux* locus and a locus within PNH region located further from the peri-nucleolar region indicated by NCL staining in the CX-5461-treated cells (Fig.4m). Interestingly, by performing the same analysis as above on public Hi-C data of mouse pre-implantation embryo development^21^, we observed that a similar trend of 3D chromatin structure reorganization of PNH and *Dux* regions when comparing early 2-cell embryos with ICM stage embryos, namely, the 2-cell embryos showed less organized PNH and less contacts between the *Dux* Locus and the PNH (Extended Data Fig.S4a-S4d), suggesting a process of maturation of PNH 3D structure organization and of the *Dux* release from the PNH during embryo development. Interestingly, we observed that 3D structure reorganization of PNH was initiated during early 2-cell to late 2-cell transition, which coincides with the beginning of shutting down *Dux* gene expression during murine early embryo development (Extended Data Fig.S4a-S4d and Extended Data Fig.S6d).

**Fig.4:**
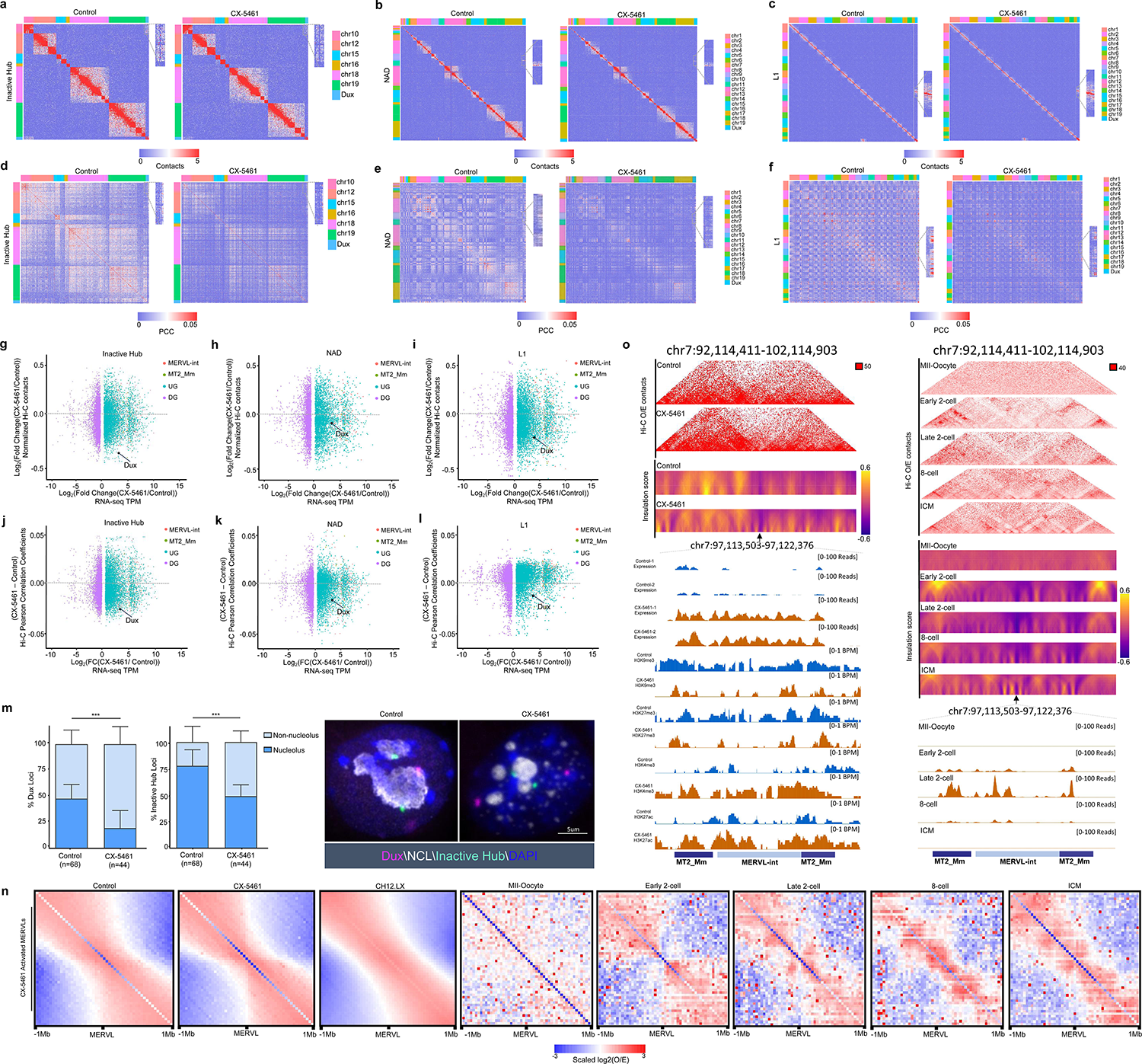
rRNA biogenesis defect drove 3D chromatin structure reorganization of PNH and *MERVL* regions towards the 2C-like state. **a)** Hi-C contact maps of Inactive Hub and 1.5 Mb genomic regions around *Dux* at 150kb resolution (GEO accession GSE166041). **b)** Hi-C contact maps of NAD and 1.5 Mb genomic regions around *Dux* at 150kb resolution (GEO accession GSE166041). **c)** Hi-C contact maps of L1 and 1.5 Mb genomic regions around *Dux* at 150kb resolution. The zoomed-in regions aim to demonstrate the change of Hi-C contacts between *Dux* and chromosome 10 in control mES cells and CX-5461 treated mES cells (GEO accession GSE166041). **d)** Hi-C pearson correlation coefficient (PCC) heat maps of Inactive Hub and 1.5Mb genomic regions around *Dux* at 150kb resolution (GEO accession GSE166041). **e)** Hi-C pearson correlation heat maps of NAD and 1.5Mb genomic regions around *Dux* at 150kb resolution (GEO accession GSE166041). **f)** Hi-C pearson correlation heat maps of L1 and 1.5Mb genomic regions around *Dux* at 150kb resolution. The zoomed-in regions aim to demonstrate the change of Hi-C PCC between *Dux* and chromosome 10 in control mES cells and CX-5461 treated mES cells (GEO accession GSE166041). **g)** Scatter plot demonstrates the log_2_(fold change) of Hi-C contacts between Inactive Hub and different types of genes in control and CX-5461 treated mES cells (GEO accession GSE166041). **h)** Scatter plot demonstrates the log_2_(fold change) of Hi-C contacts between NAD and different types of genes in control and CX-5461 treated mES cells (GEO accession GSE166041). **i)** Scatter plot demonstrates the log_2_(fold change) of Hi-C contacts between L1 and different types of genes in control and CX-5461 treated mES cells (GEO accession GSE166041). **j)** Scatter plot demonstrates the PCC difference between Inactive Hub and different types of genes in control and CX-5461 treated mES cells (GEO accession GSE166041). **k)** Scatter plot demonstrates the PCC difference between NAD and different types of genes in control and CX-5461 treated mES cells (GEO accession GSE166041). **l)** Scatter plot demonstrates the PCC difference between L1 and different types of genes in control and CX-5461 treated mES cells. The difference of PCC is defined as the average (PCC) of Inactive Hub regions and different types of genes in CX-5461 treated mES cells minus the average PCC of Inactive Hub regions and different types of genes in wild type mES cells. PCC: Pearson Correlation Coefficient, *MERVL-int*: Up-regulated *MERVL-int* genes, *MT2_Mm*: Up-regulated *MT2_Mm* genes, UG: Up-regulated genes, DG: Down-regulated genes (GEO accession GSE166041). **m)** DNA FISH analysis with a *Dux* locus probe and Inactive Hub locus probe, and co-immunostained with NCL protein. The percentage of Nucleolus-localized (overlapped with NCL) and Nucleoplasm-localized (nonoverlapped with NCL) of FISH signals is calculated. ***: p<0.001, two-way ANOVA, error bar: standard error of the mean, n denotes the number of observed mES cells (GEO accession GSE166041). **n)** Aggregate Observed(O)/Expected(E) Hi-C matrices centered on CX-5461 induced *MERVL* genes in control,CX-5461 treated mES cells (GEO accession GSE166041) and lymphoblastoid cells (GEO accession GSE63525) and mouse embryos throughout mouse pre-implantation embryonic development (GEO accession GSE82185). **o)** Representative 40kb Hi-C O/E interaction matrices of a *MERVL* loci located at TAD boundaries (chr7:97,113,503-97,122,376) are shown as heatmaps, along with the insulation score and genome browser tracks of RNA-Seq, H3K9me3, H3K27me3, H3K4me3 and H3K27ac ChIP-Seq signals of the expanded genomic region containing the TAD boundary (arrows) in control and CX-5461 treated mES cells as well as in mouse early embryos (GEO accession GSE166041 and GSE82185).

As it has been reported that 2CLCs display increased three-dimensional structural plasticity relative to mES cells^23^, we next asked whether the global 3D chromatin architecture is changed in CX-5461-treated mES cells. We compared control and CX-5461-treated mES cells Hi-C maps, with lymphoblastoid cells as a reference for full differentiated cells^59^. A global analysis of A(active)/B(inactive) compartment strength showed a slight decrease of contacts within the B compartments in CX-5461-treated mES cells compared with control mES cells (Extended Data Fig.S4e-S4h). However, at topologically associating domain (TAD) or chromatin loop level, we found a mild increase in their strength in CX-5461-treated mES cells (Extended Data Fig.S4i-S4m). To specifically investigate 2C-related genes, we further performed an analysis of local architectural differences around *MERVL* loci. We found that the insulation scores^60^ of chromatin around *MERVL* genes activated by CX-5461 treatment is markedly increased both globally (Extended Data Fig.S4n) and at local *MERVL* sites (Fig.4o), similar as observed in 2-cell embryos compared with ICM^23^ (Fig.4o), and the topological associated domain (TAD) structure around *MERVL* gene loci is more obvious (Fig.4o and Extended Data Fig.S4o), showing a more similar pattern as that in 2-cell embryos^21^ (Fig.4n). The chromatin structure reorganization around *MERVL*s is accompanied with more open chromatin states around *MERVLs* both globally (Extended Data Fig.S4n) and locally (Fig.4o and Extended Data Fig.S4o) and with their increased expression (Extended Data Fig.S4n). These results together demonstrate that nucleolar stress promoted the transformation of mES cells to 2C-like cells with reshaped 3D chromatin structure and its associated epigenetic status to facilitate gene expression, particularly at the PNH and *MERVL* regions.

### Genetic perturbation of rRNA biogenesis recapitulated CX-5461-induced 2C-like molecular phenotypes

To further investigate the critical role of rRNA biogenesis in regulating the 2C program and the homeostasis between mES cells and 2C-like cells, we generated two rRNA biogenesis-inhibited mES cell lines: 1) a line with degraded Pol I protein (PRA1) by an auxin-inducible degron system^61^ (Fig S5a), and 2) a snoRNA knockout line (SNORD113-114 gene cluster, a gift from Pengxu Qian) (Fig S5b), as snoRNAs are required for rRNA modification and biogenesis. Using these two cell lines, we performed DNA FISH experiments and found the similar molecular phenotypes of CX-5461-treated mES cells, i.e., the *Dux* locus and a representative locus within PNH region located further from the peri-nucleolar region (Fig.5a-5d). We then carried out FRAP experiments and consistently observed significantly increased mobility of NCL and NPM1 proteins in these two cell lines compared with their wild-type controls (Fig.5e-5h and Extended Data Fig.S5c-S5f). In line with these, we found that the 2C marker genes, including *MERVL*, *Dux*, *Zscan4d*, *Gm12794* and *Gm4340*, were significantly activated in these two cell lines (Fig.5i and Fig.5j). Moreover, using FACS analysis, we observed a significant increase of the percentage of tdTomato positive cells in these two cell lines compared with control cells (Fig.5k, Fig.5l, Extended Data Fig.S5g and Extended Data Fig.S5h). Collectively, these results further confirmed that repressing rRNA biogenesis can activate 2C-like program and induce the transition of mES cells to 2C-like cells.

**Fig.5:**
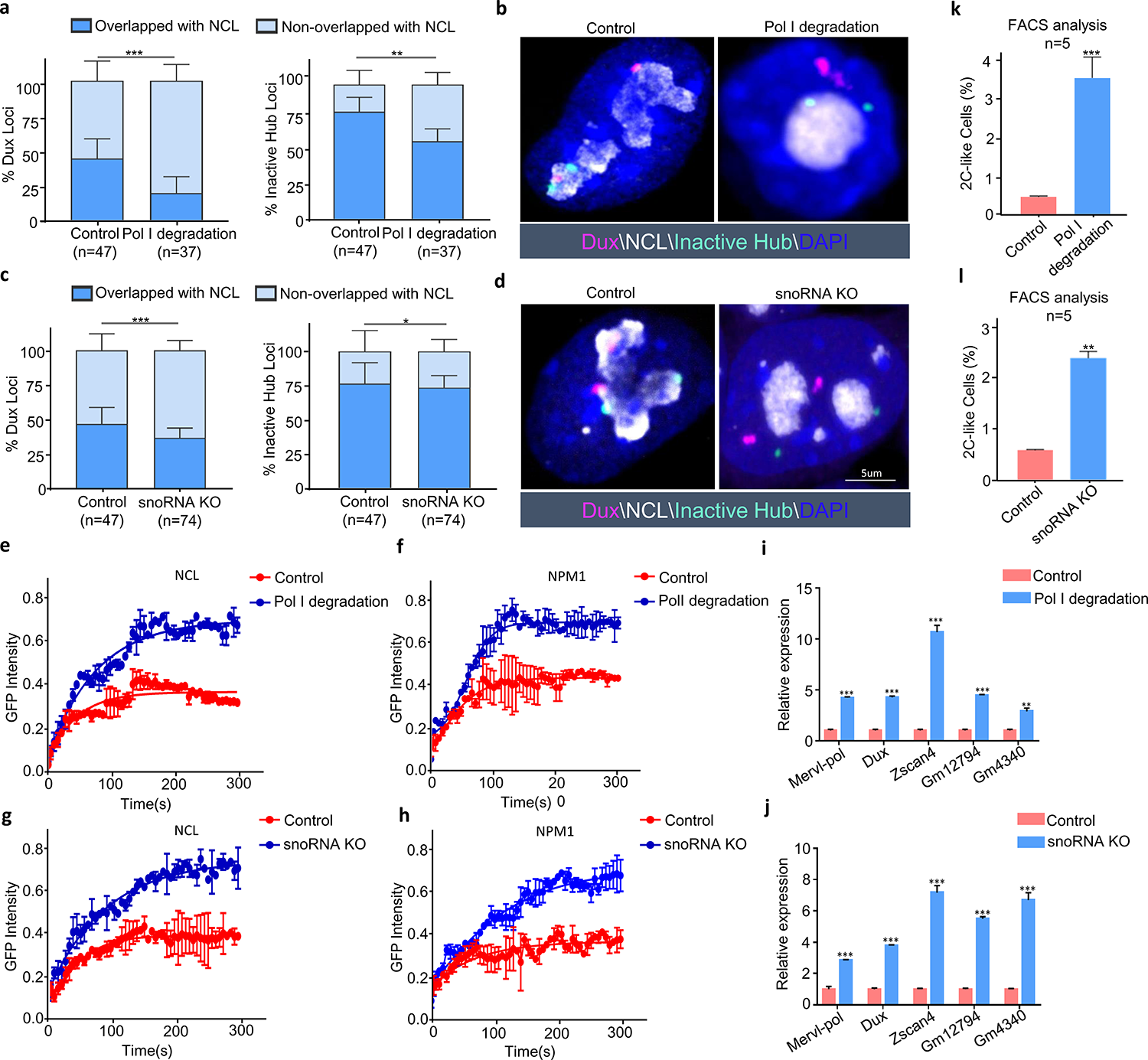
Genetic interferences of rRNA biogenesis recapitulate CX-5461-induced 2C-like molecular phenotypes. **a)** DNA FISH analysis with a *Dux* locus probe and Inactive Hub locus probe, and co-immunostained with NCL protein in control and Pol I degradation mES cell lines. **b)** The percentage of Nucleolus-localized (overlapped with NCL) and Nucleoplasm-localized (nonoverlapped with NCL) of FISH signals in control and Pol I degradation mES cell lines; **: p<0.01, ***: p<0.001, two-way ANOVA, error bar: standard error of the mean, n denotes the number of observed mES cells. **c)** DNA FISH analysis with a *Dux* locus probe and Inactive Hub locus probe, and co-immunostained with NCL protein in control and snoRNA knockout mES cell lines. **d)** The percentage of Nucleolus-localized (overlapped with NCL) and Nucleoplasm-localized (nonoverlapped with NCL) of FISH signals in control and snoRNA knockout mES cell lines; *: p<0.05, ***: p<0.001, two-way ANOVA, error bar: standard error of the mean, n denotes the number of observed mES cells. **e)** FRAP analysis showing Pol I degradation causes accelerated recovery after photobleaching of NCL, the replicates of experiment n = 4. **f)** FRAP analysis showing Pol I degradation causes accelerated recovery after photobleaching of NPM1, the replicates of experiment n = 4. **g)** FRAP analysis showing snoRNA knockout causes accelerated recovery after photobleaching of NCL, the replicates of experiment n = 4. **h)** FRAP analysis showing snoRNA knockout causes accelerated recovery after photobleaching of NPM1, the replicates of experiment n = 4. **i)** qRT-PCR quantification of 2C marker gene expression in control mES cells and Pol I degraded mES cell lines; **: p<0.01, ***: p<0.001, two-way ANOVA, the replicates of experiment n=3; error bar: standard error of the mean. **j)** qRT-PCR quantification of 2C marker gene expression in control mES cells and snoRNA knockout mES cell lines; **: p<0.01, ***: p<0.001, two-way ANOVA, the replicates of experiment n=3; error bar: standard error of the mean. **k)** The percentage of 2C::tdTomato positive cells was quantified using FACS analysis in control mES cells and Pol I degraded mES cells; Data are means ± SD, SD: Standard Deviation, ***: p<0.001, two-way ANOVA, the replicates of experiment n=5. **l)** The percentage of 2C::tdTomato positive cells was quantified using FACS analysis in control mES cells and snoRNA knockout mES cells; Data are means ± SD, SD: Standard Deviation, **: p<0.01, two-way ANOVA, the replicates of experiment n=5.

### rRNA biogenesis is critically required at the 2-cell-to-4-cell stage transition during pre-implantation embryo development

To better understand whether the physiological function of rRNA biogenesis is to facilitate embryo development during and after the 2-cell exit, we first inspected rRNA expression levels in all stages of pre-implantation mouse embryos. The precursor and matured rRNA levels were low in MII-oocyte, pronuclear zygote, early 2-cell and middle 2-cell stages but increased sharply from the late 2-cell stage to the blastocyst stage (Fig.6a). We then analyzed the expression levels of different subunit genes of RNA polymerase I (Pol I) in pre-implantation embryos. Different from rRNA expression, Pol I gene mRNA levels increased significantly from the late 2-cell stage to 4-cell stage but decreased markedly as embryos progressed through 8-cell, morula stages, and reached to blastocysts which still had a higher level than those stages before the late 2-cell (Fig.6b). Consistent with Pol I genes, we observed the similar pattern of increased expression of ribosome biogenesis genes (Extended Data Fig.S6a). This indicated that while the levels of rRNA was gradually accumulated during pre-implantation embryo development, the rRNA biogenesis rate reached to a peak during the late 2-cell to the 4-cell stage. In contrast, the ERV and 2C marker genes, such as *Dux*, *Zscan4d* and *Gm12794* were significantly decreased during the late 2-cell-to-4-cell stage (Extended Data Fig.S6b-S6d). This reciprocal expression pattern between 2C marker genes and rRNA biogenesis genes suggested that rRNA biogenesis may play a key role in shutting down the 2C program, as revealed by our 2CLC emergence analysis in cultured mES cells, and in promoting the transition from the 2-cell to the 4-cell stage. We next applied CX-5461 (an embryo tolerable concentration) to mouse early embryos as they progress through pronuclear zygotes to blastocysts. When compared with the control embryos, we found that CX-5461-treated embryos were indeed blocked before the 4-cell stage (Extended Data Fig.S6e). We further divided mouse embryos into four groups according to the different stages of CX-5461 treatment, including transitions of zygote-to-2-cell, 2-cell-to-4-cell, and morula-to-blastocyst, respectively (Fig.6c). Compared with the untreated control, we found that blastocyst formation rates of all three CX-5461-treated groups were decreased, and the 2-cell-to-4-cell-treated group showed the strongest decrease of the blastocyst formation rate at both early and late blastocyst stages (Fig.6d). This is consistent with the pattern of Pol I gene RNA expression and the pattern of PNH reshaping after early 2-cell stage during pre-implantation embryo development (Extended Data Fig.S4a-S4d), indicating that rRNA biogenesis is most critically required during the 2-cell-to-4-cell transition compared with other stages. In well support with this, the knockout of *Polr1a* gene led to mouse embryos arrested at 2-cell ^62^. Moreover, upon successful inhibition of rRNA biogenesis at the morula/blastocyst stage (Fig.6e), we found the disappearance of the NPM1- and NCL-marked “ring” structure and abnormal localization and reduced signal density of FBL in the nucleolus (Extended Data Fig.S6f). Importantly, we observed increased expression of 2C genes such as *Zscan4d*, *Gm4340*, *Dux* and *Mervl-pol* (Fig.6f). RNA-seq also demonstrated up-regulated C1 and C2 clusters of 2C genes (Fig 6g) defined before (Fig S1e) as well as ERV genes (Fig 6h-6j) in CX-5461-treated embryos compared with controls, consistent with the results from mES cells described above (Fig.1c and Extended Data Fig.S1d). Altogether, these data demonstrated rRNA biogenesis and nucleolar integrity is a molecular switch for the transition from the 2-cell to the 4-cell embryos.

**Fig.6:**
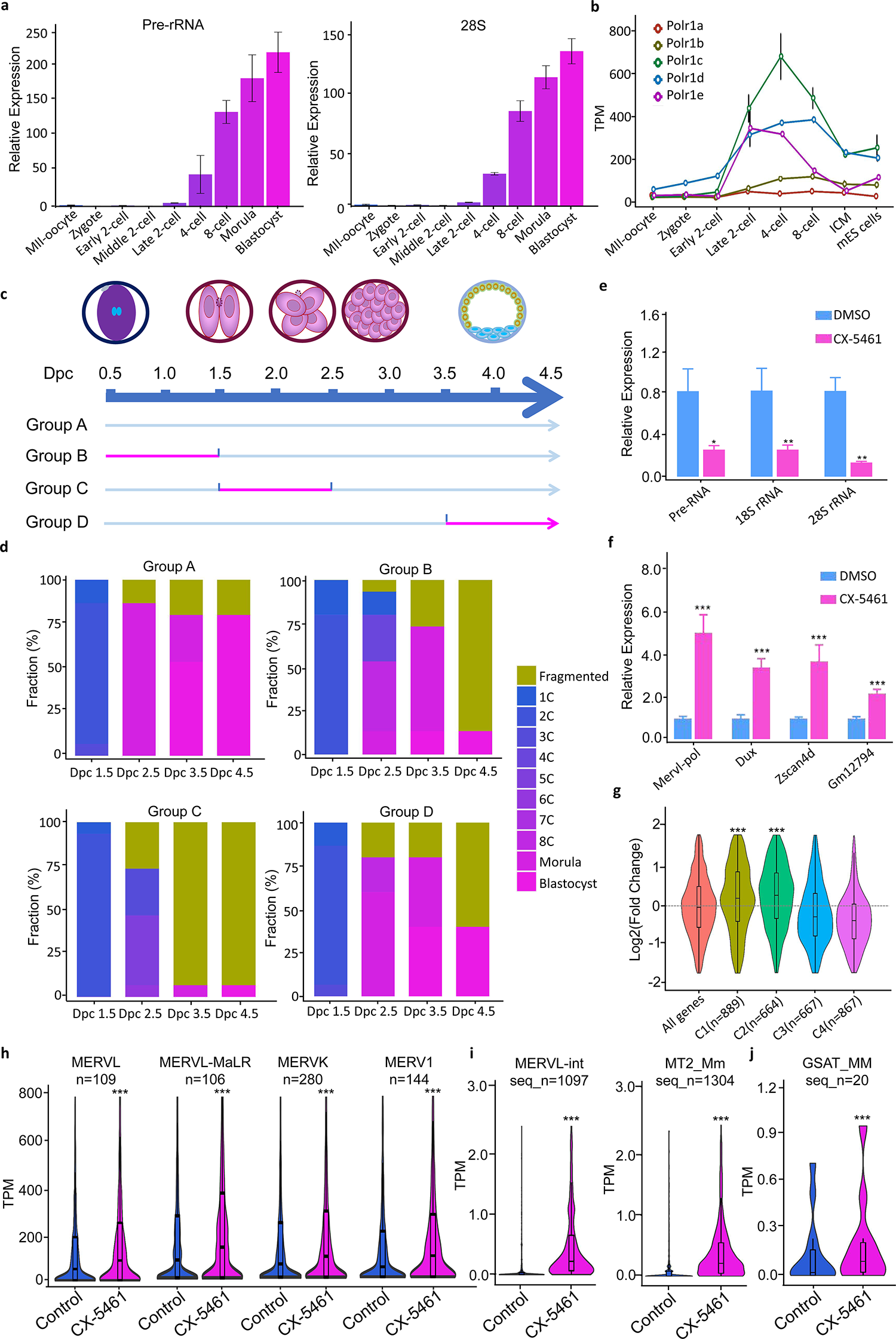
rRNA biogenesis is critically required at the 2-cell-to-4-cell stage transition during pre-implantation embryo development. **a)** Expression of pre-rRNA and 28S rRNA across different embryo developmental stages. **b)** Expression of different subunit genes of RNA polymerase I across different embryo developmental stages. **c)** Different schemes of treatment with CX-5461. The 24hrs time window for CX-5461 treatment is highlighted in red; Dpc: Days post-coitum. **d)** Stacked bar plots showing fraction of embryos at different developmental stages with the different CX-5461 treatment schemes in Fig 6c. The numbers of embryos of group A to group D were all 15 embryos. **e)** qRT-PCR showing rRNA expression level in blastocysts, after CX-5461 treatment of morula embryos followed by *in vitro* culture of the treated embryos. *: p<0.05, **: p<0.01, two-way ANOVA, the replicates of experiment n=3. **f)** qRT-PCR showing 2C marker gene expression level in blastocysts, after CX-5461 treatment of morula embryos followed by *in vitro* culture of the treated embryos. ***: p<0.001, two-way ANOVA, the replicates of experiment n=3. **g)** Violin plots demonstrating the expression level changes of stage-specific gene clusters of mouse pre-implantation embryos (as defined in Extended Data Fig S1) in CX-5461-treated and control blastocyst embryos, ***: p<0.001, Mann-Whitney U test (GEO accession GSE166041). **h)** Violin plots show expression levels of major ERV gene classes in control blastocyst embryos and CX-5461 treated blastocyst embryos; n denotes the number of sub-classes of ERV genes; ***: p<0.001, Wilcox signed rank test (GEO accession GSE166041). **i)** Violin plots show expression levels of ERV gene sub-classes of *MERVL-int* and *MT2_Mm* in control blastocyst embryos and CX-5461 treated blastocyst embryos; seq_n denotes the number of annotated *MERVL-int* and *MT2_Mm* sequences in the mouse mm10 reference genome; *: ***: p<0.001, Wilcox signed rank test (GEO accession GSE166041). **j)** Violin plots show expression levels of ERV gene sub-classes of *GSAT_MM* in control blastocyst embryos and CX-5461-treated blastocyst embryos; seq_n denotes the number of annotated *GSAT_MM* sequences in the mouse mm10 reference genome; ***: p<0.001, Wilcox signed rank test (GEO accession GSE166041).

## Discussion

Starting from zygotic genome activation (ZGA) at the 2-cell stage, an embryo undergoes through the four/eight-cell, morula, blastocyst stages, and then prepares itself for implantation. Along with this pre-implantation development process, basic anabolic metabolism and translational processes become more active. Nucleoli, the organelles involved in translation, functionally mature from nucleolar precursor bodies (NPB) during this process^63–65^. Interestingly, a recent research work reported that TRIM28/Nucleolin/LINE1 complex that can mediate both ZGA gene Dux repression and rRNA expression^14^, suggesting that shutting down ZGA and initiating nucleoli formation are not independent events but interconnected. However, a complete picture on how the transition between ZGA and nucleolar formation occurs in the nucleus is not fully understood.

Here, we reported that nucleolar rRNA biogenesis and higher-order 3D chromatin structure remodeling of PNH might coordinate to develop during the 2-cell to later stage transition, and we found that mES cells cultured *in vitro* can transform into 2C-like cells upon nucleolar stress caused by repressing rRNA biogenesis. We propose a mechanistic model for the novel role of rRNA biogenesis in regulating the 2C-like program and the homeostasis between 2C-like cells and mES cells (Fig 7). In the unperturbed mES cells, nucleolar integrity mediated by rRNA biogenesis maintains the normal the liquid-liquid phase separation (LLPS) of nucleolus and the formation of peri-nucleolar heterochromatin (PNH) containing *Dux*, and this normal nucleolar LLPS facilitated NCL/TRIM28 complex occupancy on the *Dux* locus to repress *Dux* expression. In contrast, in the rRNA biogenesis-repressed mES cells, the natural liquid-like phase of nucleolus is disrupted, causing dissociation of the NCL/TRIM28 complex from the PNH and changes of epigenetic state and 3D structure of the PNH, which eventually leads to *Dux* to be released from the PNH region, activation of 2C-like program and transition of mES cells to 2C-like cells. Given the dynamic regulation of nucleolus and rRNA gene chromatin during early embryo development and the sensitivity of embryos to environmental stress at the early stages, it is conceivable that embryos may use the mechanisms elucidated above to ensure its safe development.

**Fig.7:**
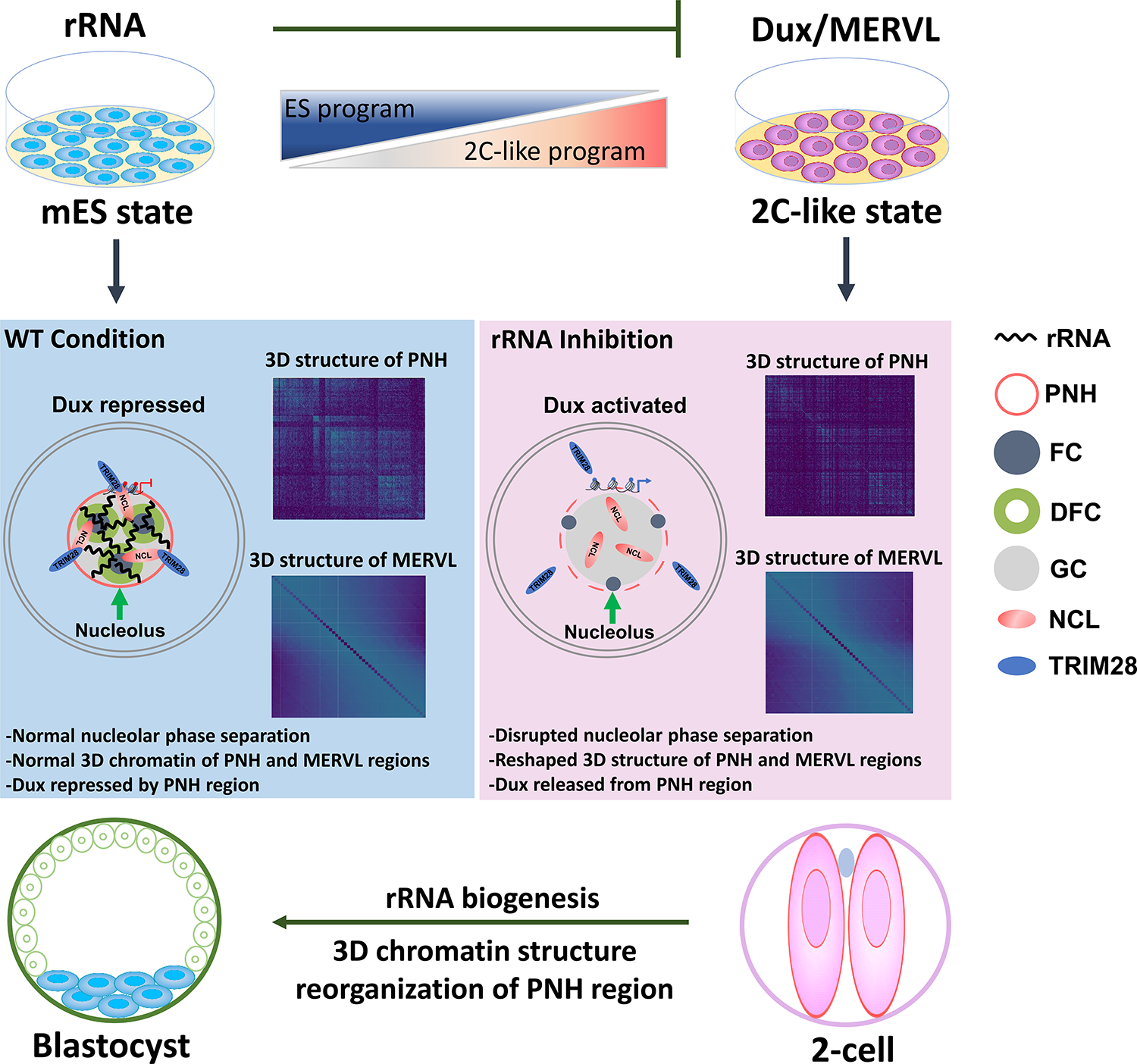
A mechanistic model for the role of rRNA biogenesis in regulating the 2C-like program and the homeostasis between 2C-like cells and mES cells. In the unperturbed mES cells, nucleolar integrity mediated by rRNA biogenesis maintains the normal the liquid-liquid phase separation (LLPS) of nucleolus and the formation of peri-nucleolar heterochromatin (PNH) containing *Dux*, and this normal nucleolar LLPS facilitated NCL/TRIM28 complex occupancy on the *Dux* locus to repress *Dux* expression. In contrast, in the rRNA biogenesis-inhibited mES cells, the natural liquid-like phase of nucleolus is disrupted, causing dissociation of the NCL/TRIM28 complex from the PNH and changes of epigenetic state and 3D structure of the PNH, which eventually leads to *Dux* to be released from the PNH, activation of 2C-like program and transition of mES cells to 2C-like cells.

Nucleolus, the largest membrane-less condensate in a cell, is a stress-sensitive organelle and ensure quality control of nuclear proteome under stress^37, 38^. Its association with heterochromatin in its periphery confers genetic regulation of key cell fate decision factors such as *Dux* in pluripotent stem cells. Previous studies on nucleolus in stem cells mainly focused on the role of rRNA and its associated chromatin in the context of ES cell self-renewal and differentiation or exit of pluripotency^44, 57, 66^. In contrast, our work for the first time provided a novel perspective in reprogramming mES cells back to 2C-like cell, and in nucleolar phase separation and 3D chromatin structure remodeling at the PNH. These findings are in line with the emerging notion that phase-separated condensates regulate transcription, epigenetics, and higher-order chromatin structure^39–41, 67–70^, and shed light on a previously neglected area of nucleolus-associated condensates in chromatin control during early development. It is being worth mentioned that we do not intend to overstate that the fate transition of mES to 2C-like cell triggered by rRNA biogenesis defect are explained by LLPS^71^. What we observed is that the integrity of nucleolus mediated by rRNA biogenesis maintains the normal nucleolar LLPS and 3D structure of PNH. It is possible that 3D structure reshaping of PNH mediated by nucleolar LLPS is a common thread of RNA and protein-mediated Dux silences and 2C repression (e.g., rRNA, snoRNA, LINE1 RNA, NCL, TRIM28 and LIN28). The NCL/TRIM28 complex or other nucleolar proteins of GC layer and heterochromatin proteins, e.g., NPM1 and HP1, is possible the key regulating factors connecting nucleolar LLPS and the establishment and maintenance of PNH. Our study gives a starting clue that the LLPS model is important for the assembly and function of nucleolus^37^ with the implication of gene regulation, and the quantitative and mechanistic models merit future investigations.

## Methods

### Cell culture

E14 mES cells were cultured on 0.1% gelatin-coated plates with MEF feeder cells in N2/B27/LIF/2i medium (1:1 mix of DMEM/F12 (11320-033, Gibco) and Neurobasal medium (21103-049, Gibco) containing 1×N2 and B27 supplements (17502-048/17504-044, Life Technologies), 100 µM non-essential amino acids (GNM71450, GENOM), and 1,000 U/ml LIF (PEPRO TECH), 1 μM PD03259010 and 3 μM CHIR99021 (STEMCELL Technologies) and 100 U/ml penicillin, 100 µg/ml streptomycin (15140-122, Gibco). For primed state media, 20 ng/ml Activin, 10 ng/ml FGF2, and 1% KSR were added to the 1:1 DMEM/F12 and Neurobasal medium containing N2 and B27. To investigate DUX binding, an N-terminal FLAG-DUX protein was expressed in our clonal cell lines. In Control group, mES cells were treated with doxycycline for 12h to induce FLAG-DUX expression and then treatment of negative-control Silencer Select siRNA. In siDux group, mES cells were treated with doxycycline for 12h and then siDUX for two days. In CX-5461 treatment group, mES cells were treated with doxycycline for 12h and then treatment of CX5461. In siDux+CX-5461 group, mES cells were treated with doxycycline for 12h and then siDUX for two days followed by treatment of CX-5461.

### Fluorescence activated cell sorting (FACS) analysis

The E14 2C::*tdTomato* cells were generated as in ^1^. E14 mES cells were transfected with 2C::*tdTomato* using Lipofectamine 2000 and selected with 150 μg/ml hygromycin 48 hr after transfection and for 7 days. Mouse E14 wild-type cells were subjected to 0.4μM CX-5461 treatment for 12h or 2μM CX-5461 treatment for 12h. Cells were isolated by FACS to measure the ratio of 2C-like cells. Apoptosis was measured using the Annexin V and DAPI Staining.

### Cell line immunofluorescence staining

E14 mES cells were grown on gelatin-coated glass coverslips with MEFs and cultured 12 h before fixed with 4% PFA for 10 min, and then permeabilized with 0.5% Triton X-100 in PBS for 20 min at room temperature (RT). The cell samples were blocked in blocking buffer (3% BSA, 2% donkey serum in PBS) for 10 min at RT and then stained with a primary antibody (1:200, Nucleolin, CST, 145745; 1:400, NPM1, Sigma, B0556; 1:200, Fibrillarin, Abcam, ab4566; RPA194, 1:50, Santa Cruz, sc-48385) for 12 h at 4 °C. After three washes with 0.1% Triton X-100 in PBS, cells were stained with a secondary antibody (1:200, Goat polyclonal Secondary Antibody to Mouse IgG, Abcam, ab150113) for 2-12 h at 4 °C. Followed by washing three times with 0.5% Triton X-100/PBS, DAPI was used for nucleus staining. The samples were then imaged by Zeiss LSM880 fluorescence microscope at a 63×oil objective. For high regulation microscopy imaging, LSM800 with Airyscan module was used.

### Fluorescence recovery after photobleaching (FRAP) analysis

E14 wild-type and CX-5461 treatment mES cells cultured on MEF cells were grown in N2B27/LIF/2i conditions and maintained at 37 °C and with 5% CO2 during image acquisition. Cells were transduced with Lenti-NCL-GFP / Lenti-NPM1-GFP / Lenti-FBL-mCherry lentivirus. FRAP experiments were performed on a ZEISS (Jena, Germany) LSM800 confocal laser scanning microscope equipped with a ZEISS Plan-APO 63x/NA1.46 oil immersion objective. Circular regions of constant size were bleached and monitored overtime for fluorescence recovery. Bleaching was once every 10 seconds for a total of 10 minutes. Fluorescence intensity data was corrected for background fluorescence and normalized to initial intensity before bleaching using GraphPad software. Resulting FRAP curves were fitted with Four parameter logistic (4PL) curve.

### siRNA-mediated knockdown in mES cells

siRNA transfections were performed in mES cells with Lipofectamine 2000 (Thermo Fisher Scientific). mES cells were seeded into 12-well plate and cultured in LIF/2i medium for overnight. The next day, 800 μl LIF/2i medium without antibiotics was added into each well. Then, the transfect mixture (40 pmol of 3 independent siRNA targeting each gene/a non-targeting siRNA (negative control, NC) and 2 μl of Lipo 2000 which was diluted in 200 μl Opti-MEM medium (Gibco)) was added into each well and incubated for 6 hr at 37 °C. After incubation, the medium was exchanged for fresh complete LIF/2i medium and cells were harvested for RNA extraction approximately 48 hr later. The sequences of siRNA are listed in Supplementary Table 1.

### Cell line RNA extraction and qRT-PCR

Total RNA was isolated from mES cells using miRNeasy kit (217004, QIAGEN) according to the manufacturer’s protocol, and 1 μg RNA was reverse transcribed to cDNA with HiScript II Q RT Super Mix (R223-01, Vazyme). Gene expression was analyzed with SYBR-Green qPCR Master mix (Bio-Rad) on Bio-Rad PCR machine (CFX-96 Touch). Each gene was normalized to Actin or Gapdh. All primers used are listed in Supplementary Table 2.

### Oligopaint DNA FISH

The Oligopaint DNA FISH probe was designed according to several previous publication. The encoding probes were designed as previous described^72, 73^ and the targeted genomic region was designed by OligoMiner^74^. The probe pools synthesized by Synbio Technologies were used as templates and the dye-labeled secondary probes were produced by Sunya Biotechnology. All sequences used in this work were listed in Supplementary Table 3. Briefly, the synthesized probe pool was first used as template to amplify via 30 PCR cycles and was subsequently purified by ammonium acetate precipitation. Then, the previous PCR products was used as template to amplify and convert into RNA via an *in vitro* transcription of high yield (New England Biolabs, E2040S); the RNA product above were converted back into single-stranded DNA via reverse transcription. At last, the product was subjected to the alkaline hydrolysis to remove template RNA and was further purified by ammonium acetate precipitation. For the secondary probe, a 30 bp random oligo was designed as previous and attached with Cy3 or Alexa Flour 647 at 5’ end.

For DNA FISH, the cell samples were fixed in 4% Paraformaldehyde (Sigma, 158127) for 10 minutes and washed two times with PBS, followed by permeabilizing with 0.5% Triton X-100 (Sigma, T8787) in PBS. Then the samples were incubated in 0.1% w/v sodium borohydride (Sigma, 71320) for 10 minutes and treated with 0.1M HCl for 5 minutes. After that, the cell samples were incubated in 0.1 mg/ml RNase A diluted in PBS for 45 minutes in 37°C. After washing three times in 2x SSCT (2x SSC + 0.1% Tween-20), the samples were immersed in 50% formamide diluted in 2x SSCT for 15 minutes at room temperature then transferred to 85°C for 10 minutes. The primary probe and secondary probe were freshly mixed into hybridization buffer (2x SSC, 50% formamide, 20% dextran sulfate) at 6 µM and 1 µM final concentration and dropped on samples. The samples were heated at 85 °C for 20 minutes and transferred to a 37 °C incubator before hybridization overnight. For the co-immunostaining, the cell samples above were washed three times in PBS and then incubated with primary antibody (1:200, Nucleolin, CST, 145745) for 12 h at 4°C. After three times of washing with PBS, Donkey anti-Rabbit secondary antibody (1:200, Abcam, ab150077) was performed and incubated for 4 h. After washing three times with PBS, DAPI was used for nucleus staining.

### Transmission electron microscope (TEM)

For transmission electron microscopy, E14 mES cells cultured in one 10 cm dish were collected, cleaned from feeder cells, and supplemented with 2.5% glutaraldehyde. The cell pellet was dispersed into small clusters and fixed at least 6 hr at 4 °C. After that, the cell samples were treated with standard procedures. Then the slices were imaged on FEI Spirit 120 kV LaB6 Routine Cryo-EM Capable Electron Microscope.

### Embryo collection and culture

The C57BL/6J mice were housed in the animal facility of Zhejiang University. All experimental procedures were performed in accordance with the Animal Research Committee guidelines of Zhejiang University. To collect pre-implantation embryos, C57BL/6J female mice (4–6 weeks old) were intraperitoneally injected with 7.5 IU each of PMSG (San-Sheng Pharmaceutical) for 48h followed by injection of 7.5 IU of hCG (San-Sheng Pharmaceutical). The superovulated female mice were mated with adult males overnight after hCG administration. Embryos at different stages of pre-implantation development were collected at defined time periods after the administration of hCG: 30 h (early 2-cell), 44–48 h (2-cell), 54–56 h (4-cell), 68–70 h (8-cell), 76-78 h (morula) and 92–94 h (blastocysts) in HEPES-buffered CZB medium. Zygotes were collected from ampullae of oviducts and released with hyaluronidase for removing cumulus cells.

### Embryo immunofluorescence staining

Embryos were first fixed with 1% and 2% paraformaldehyde (PFA) in 1×PBS for 3 min sequentially, followed by treatment with 4% PFA for 30 min at room temperature (RT). Embryos were washed three times with 1×PBS, permeabilized for 15 minutes in PBS/0.25% Triton X-100 and blocked-in blocking buffer (PBS/0.2% BSA/0.01% Tween-20) for 1 hr at RT, followed by incubation overnight with primary antibodies (1:200, Nucleolin, CST, 145745; 1:400, NPM1, Sigma, B0556; 1:200, Fibrillarin, Abcam, ab4566; RPA194, 1:50, Santa Cruz, sc-48385) at 4 degree or for 1hr at 37 °C. Subsequently, embryos were washed four times for 10 min each and incubated with a secondary antibody (daylight 488-conjugated anti-rabbit, 1:100 or daylight 594-conjugated anti-mouse, 1:200) for 1hr at 37°C and washed three times with PBS. Nuclei were stained with DAPI for 1 min. Embryos were observed under Zeiss LSM880 fluorescence microscope at 63× magnification with an oil immersion objective.

### Embryo collection, cDNA synthesis and qRT-PCR

10 embryos were rinsed in 0.2% BSA/PBS without Ca2+ and Mg2+ and placed in 0.2 ml PCR tube, immediately transferred in liquid nitrogen, and stored at -80 °C. It was hybridized with 0.5 µl oligo-dT30 (10 μM, Takara) and 1 μl random (1 M) and 1 µl dNTP mix (10 mM) in 2 μl cell lysis buffer (2 U RNase inhibitor, 0.01% Triton X-100) at 72 °C for 3 min. Then, the reaction was immediately quenched on ice. After the reaction tube was centrifuged, 2 µl was used for reverse transcription with Super Script II Reverse Transcriptase 5x first strand buffer, 0.25 µl RNase inhibitor (40 U), 0.06 µl MgCl2 (1 M), 2 µl betaine (5 M), and 0.5 µl Reverse Transcriptase Superscript II (Takara). Reverse transcription was carried out in the thermocycler at 42 °C for 90 min, 70 °C for 15 min, and then 4 °C for holding. Subsequently, cDNA was diluted 1:10 (v/v) with RNase free water and used for a qPCR amplification in triplicate with SYBR Green Master (Vazyme) in a final volume of 20 µl per reaction as manufacturer’s instructions.

### Embryo RNA-seq library preparation and sequencing

Embryos were collected (5 embryos per sample) in 0.2 ml PCR tubes with a micro-capillary pipette and processed into cDNA with Superscript II reverse transcriptase. The cDNA is amplified with KAPA Hifi HotStart using 12 cycles. Sequencing libraries were constructed from 1 ng of pre-amplified cDNA using DNA library preparation kit (TruePrep DNA Library Prep Kit V2 for Illumina, Vazyme). Libraries were sequenced on a HiSeq-PE150, with paired end reads of 150 bp length each.

### Bulk RNA-seq library preparation and sequencing

A total amount of 2 μg RNA per sample was used as input materials for the RNA sample preparation. mRNA was purified from total RNA using poly-T oligo-attached magnetic beads. Purified mRNA was fragmented at 94 °C for 15 min by using divalent cations under elevated temperature in NEBNext first strand synthesis reaction buffer (5X). First strand cDNA was synthesized using random primer and ProtoScript II reverse transcriptase in a preheated thermal cycler as follows: 10 min at 25 °C; 15 min at 42 °C; 15 min at 70 °C. Immediately finished, second strand synthesis reaction was performed by using second strand synthesis reaction buffer (10X) and enzyme mix at 16 °C for 1 hr. The library fragments were purified with QiaQuick PCR kits and elution with EB buffer, then terminal repair, A-tailing and adapter added were implemented. The products were retrieved, and PCR was performed for library enrichment. The libraries were sequenced on an Illumina platform.

### 10X single-cell mRNA library preparation and sequencing

Single-cell suspensions of control and CX-5461 treated mES cells were resuspended in DPBS-0.04% BSA at 1x106 cells/mL. Then scRNA-seq libraries were generated from the 10X Single Cell 3’ Solution Reagents V2 according to the manufacturer’s protocol (10x Genomics). After the GEM-RT incubation, barcoded-cDNA was purified with DynaBeads cleanup mix, followed by 10-cycles of PCR amplification (98°C for 3 min; [98°C for 15 s, 67°C for 20 s, 72°C for 1 min] x 10; 72°C for 1 min). The total cDNA of single-cell transcriptomes was fragmented, double-size selected with SPRI beads (Beckman), followed by 12 cycles sample index PCR amplification (98°C for 45 s; [98°C for 20 s, 54°C for 30 s, 72°C for 1 min] x 10; 72°C for 1 min), then another double-size selection with SPRI beads was performed before sequencing. Libraries were sequenced on the Illumina Hiseq X10 platform according to the manufacturer’s instructions (Illumina). Read 1 and Read 2 (paired end) were 150 bp, and the length of index primer was designed as 8 bp.

### ChIP-seq library preparation and sequencing

mES cells were cross-linked in 1% formaldehyde for 10 min at 37 °C, followed by adding glycine to a final concentration of 125 mM and incubated for 5 min at room temperature. Spin the cells for 5 min at 4 °C, 1100 rpm, and wash twice in ice-cord PBS. Cell pellet was resuspended with lysis buffer containing 1× Protease Inhibitor Cocktail and incubated on ice for 10 min, then vortexed vigorously for 10 seconds and centrifuged at 3000 rpm for 5 minutes. The pellet was re-suspended in ChIP lysis buffer and incubated on ice for 10 minutes and vortexed occasionally. Afterwards, the chromatin lysate was transferred to a 1.5 ml centrifuge tube and chromatin sheared using water bath sonication with the following conditions: shear 15 cycles at 4 °C, 15 seconds on, 30 seconds off. Centrifuge and transfer supernatant to a new tube. Taking 5 μl (1%) from the 500 μl containing sheared chromatin as input. Each chromatin sample was incubated with antibodies for H3K9me3 Rabbit polyclonal antibody (abcam, ab8898), H3K27me3 Rabbit mAb (CST, 9733), H3K4me3 Rabbit mAb (CST, 9751), H3K27ac Rabbit mAb (CST, 8173), Nucleolin (D4C7O) Rabbit (CST, 14574), TRIM28 Mouse monoclonal (20C1) (abcam, ab22553) overnight on a rotating platform at 4 °C. The next day, the sample was incubated with protein A+G magnetic beads (HY-K0202, MCE) for 3 hr at 4 °C with rotation. The beads-antibody/chromatin complex was washed three times with low-salt wash buffer and once with high-salt wash buffer and resuspended with elution buffer. The elute DNA was treated with RNase A at 42 °C for 30 min, then treated with protease K at 60 °C for 45 min followed by heat inactivation at 95 °C for 15 min. The purified DNA was subjected to library preparation or analyzed by qPCR. The libraries were sequenced on an Illumina platform.

### *In situ* Hi-C library preparation and sequencing

10^6 cells were cross-linked for 10 min with 1% final concentration fresh formaldehyde and quenched with 0.2M final concentration glycine for 5 min. The cross-linked cells were subsequently lysed in lysis buffer (10 mM Tris-HCl (pH 8.0), 10 mM NaCl, 0.2% NP40, and complete protease inhibitors (Roche)). The extracted nuclei were re-suspended with 150 μl 0.1% SDS and incubated at 65°C for 10 min, then SDS molecules were quenched by adding 120 μl water and 30 μl 10% Triton X-100, and incubated at 37 °C for 15 min. The DNA in the nuclei was digested by adding 30 μl 10x NEB buffer 2.1 (50 mM NaCl, 10 mM Tris-HCl, 10 mM MgCl2, 100 μg/ml BSA, pH 7.9) and 150U of MboI, and incubated at 37 °C overnight. On the next day, the MboI enzyme was inactivated at 65 °C for 20 min. Next, the cohesive ends were filled in by adding 1 μl of 10 mM dTTP, 1μl of 10 mM dATP, 1 μl of 10 mM dGTP, 2 μl of 5mM biotin-14-dCTP, 14 μl water and 4 μl (40 U) Klenow, and incubated at 37 °C for 2 h. Subsequently, 663 μl water,120 μl 10x blunt-end ligation buffer (300 mM Tris-HCl, 100 mM MgCl2, 100 mM DTT, 1 mM ATP, pH 7.8), 100μl 10% Triton X-100 and 20 U T4 DNA ligase were added to start proximity ligation. The ligation reaction was placed at 16 °C for 4 h. After ligation, the cross-linking was reversed by 200 µg/mL proteinase K (Thermo) at 65°C overnight. DNA purification was achieved through QIAamp DNA Mini Kit (Qiagen) according to manufacturer’s instructions. Purified DNA was sheared to a length of ∼400 bp. Point ligation junctions were pulled down by Dynabeads® MyOne™ Streptavidin C1 (Thermofisher) according to manufacturer’s instructions. The Hi-C library for Illumina sequencing was prepped by NEBNext® Ultra™ II DNA library Prep Kit for Illumina (NEB) according to manufacturer’s instructions. The final library was sequenced on the Illumina HiSeq X Ten platform (San Diego, CA, United States) with 150PEmode. Two replicates were generated for one group material.

### Bulk RNA-seq data analysis

All bulk RNA-seq reads were trimmed using Trimmomatic software (Version 0.36) with the following settings “ILLUMINACLIP:TruSeq3-PE.fa:2:30:10 LEADING:3 TRAILING:3 SLIDINGWINDOW:4:15 MINLEN:36”^75^ and were further quality-filtered using FASTX Toolkit (http://hannonlab.cshl.edu/fastx_toolkit/) fastq_quality_trimmer command with the minimum quality score 20 and minimum percent of 80% bases that has a quality score larger than this cutoff value. The high-quality reads were mapped to the mm10 genome by HISAT2, a fast and sensitive spliced alignment program for mapping RNA-seq reads, with -dta parameter^76^. PCR duplicate reads were removed using Picard tools (https://broadinstitute.github.io/picard/). For subsequent analysis on single-copy genes, only uniquely mapped reads were kept. Considering the multi-mapping of reads derived from repeat sequences, we used all mapped reads for further analysis. The expression levels of genes and repeat sequences were independently calculated by StringTie^77^ (Version v1.3.4d) with -e -B -G parameters using Release M18 (GRCm38.p6) gene annotations downloaded from GENCODE data portal and annotated repeats (RepeatMasker) downloaded from the UCSC genome browser, respectively. To obtain reliable and cross-sample comparable expression abundance estimation for each gene and each family of repeat sequence, reads mapped to mm10 were counted as TPM (Transcripts Per Million reads) based on their genome locations. Differential expression analysis of genes in different samples was performed by DESeq2 using the reads count matrix produced from a python script “prepDE.py” provided in StringTie website (http://ccb.jhu.edu/software/stringtie/). We selected the genes with stage-specific scores larger than 0.2 to perform K-mean clustering analysis using pheatmap R package. The stage-specific scores of genes expressed during mouse early embryo development were obtained by entropy-based measure^78^.

Unsupervised hierarchical clustering was carried out to compare the transcriptomes of mES cells from our study and other reports of 2C-like cell (2C::*tdTomato*+ and 2C::*tdTomato*-(GEO accession GSE33923); Zscan4_Em+ and Zscan4_Em-(GEO accession GSE51682); Kap1_KO and Kap1_WT (GEO accession GSE74278); CAF1_KO and CAF1_WT (GEO accession GSE85632), Dux_GFPpos and Dux_GFPneg (GEO accession GSE85632); LINE1 ASO and RC ASO (GEO accession GSE100939); Dppa4_GFPpos and Dppa4_GFPneg (GEO accession GSE120953), NELFA_GFPpos and NELFA_GFPneg (GEO accession GSE113671); Lin28a_KO and Lin28a_WT (GEO accession GSE164420). Additionally, pre-implantation mouse embryos of different developmental stages (GSE66582) were included for comparison. TPM were obtained for each sample using the StringTie described above. Only genes that were expressed TPM ≥ 1 were included for analysis. A log2 transformation was applied after adding one pseudo-count (that is, log2[TPM+1]). The ComBat function from the sva package (https://bioconductor.org/packages/release/bioc/html/sva.html) was applied on log2 expression values to correct for batch effects caused by different experiments and sequencing platforms.

### Single-cell RNA-seq data analysis

For single cell RNA-seq, we used 10x Genomics system following the manufacturer’s protocol. We followed the previously published pipeline^79^ to produce digital gene expression matrices of the droplet microfluidics-based single-cell RNA-seq sequencing data derived from control and CX-5461 treated mouse ES cells. Single-cell gene expression matrix was further analyzed with Seurat (https://satijalab.org/seurat/) (v2.3.4). We excluded the genes with expressed cell number smaller than 3 and the cells with nUMIs smaller than 500 or the expression percentages of mitochondrial genes larger than 0.2 and used 16 Principle Components (PCs) for tSNE analysis. Especially, we modified the published pipeline by substituting the aligner of Bowtie with HISAT2 for calculating the expression of repeat sequences. Considering the multi-mapping of reads derived from repeat sequences, we used all mapped reads for further analysis.

### ChIP-seq and ATAC-seq data analysis

To tailor and filter ATAC-seq and ChIP-seq reads, we used the same procedure as RNA-seq reads processing. To avoid the potential effects of inconsistent sequencing depths on subsequent data analyses, we randomly sampled equal numbers read pairs from each experimental sample. For each sample, the ATAC-seq and ChIP-seq reads were first aligned to mm10 genomes using Bowtie2 (version 2.3.4.1)^80^. The ATAC-seq reads were aligned with the parameters: -t -q -N 1 -L 25 -X 2000 no-mixed no-discordant. The ChIP-seq reads were aligned to mm10 with the options: -t -q -N 1 -L 25. The ATAC-seq reads were aligned with the parameters: -t -q -N 1 -L 25 -X 2000 no-mixed no-discordant. The ChIP-seq reads were aligned to mm10 with the options: -t -q -N 1 -L 25. For meta-analysis of genome regions, all unmapped reads, multiple mapped reads, and PCR duplicates were removed. For demonstrating the sequencing signal around Dux locus in UCSC genome browser, we maintained all multiple mapped reads for visualization. The bamCoverage and bamCompare commands contained in deepTools ^81^ (version 2.5.3) were adopted for downstream analysis. Using BamCoverage command with the parameters: -normalizeUsing BPM -of bigwig -binSize 100, we normalized the raw reads signal to Bins Per Million mapped reads (BPM) signal and converted the alignment bam files to bigwig signal files. The bigwig files were imported into UCSC genome browser for visualization. To minimize the effect of chromatin structure and sequencing bias in our ChIP-seq data, we corrected ChIP-seq signal using log2 ratio transformation between H3K9me3 signal and input signal by BamCompare command. We only considered the log2 ratio larger than 0 as effective ChIP-seq signals. The “computeMatrix” and “plotProfile” commands of deepTools were used to produce the reads density distribution plot of ATAC-seq and ChIP-seq signal in the given genomic region. For meta-analysis of sequencing signals in L1 regions, we only used their subsets which have overlaps with Inactive Hub regions. Homer (v4.11)^82^ was used for motif discovery and enrichment analysis. For motifs across gene promoters, the search space was defined as a 4 kilobase (kb) window centered at the transcription start site (findMotifs.pl geneInput.txt mouse out/ -start -2000 -end 2000 -len 8,12 -p 4).

### Hi-C data analysis

The paired-end reads of fastq files were aligned, processed, and iteratively corrected using HiC-Pro (version 2.11.1) as previously described^83^. Briefly, short sequencing reads were first independently mapped to mouse mm10 reference genome using the bowtie2 aligner with end-to-end algorithm and ‘-very-sensitive’ option. To rescue the chimeric fragments spanning the ligation junction, the ligation site was detected and the 5′ fraction of the reads was aligned back to the reference genome. Unmapped reads, multiple mapped reads and singletons were then discarded. Pairs of aligned reads were then assigned to MboI restriction fragments. Read pairs from the uncut DNA, self-circle ligation and PCR artifacts were filtered out and the valid read pairs involving two different restriction fragments were used to build the contact matrix. To eliminate the possible effects on data analyses of variable sequencing depths, we randomly sampled equal numbers read pairs from each condition for downstream analyses involving comparison analyses between conditions. Valid read pairs were then binned at a 40kb, 150kb and 500 kb resolutions by dividing the genome into bins of equal size. The binned interaction matrices were then corrected by Knight–Ruiz matrix balancing method using hicCorrectMatrix command with the parameter -- correctionMethod KR in HiCExplorer (https://hicexplorer.readthedocs.io/en/latest/) (v3.3)^85^. The Observed/Expected (O/E) Hi-C contact matrix was produced by HiCExplorer hicTransform command with --method obs_exp_norm option. Pearson Correlation Coefficients Hi-C matrix was obtained by HiCExplorer hicTransform command with --method pearson option.

A and B compartments were identified using the first eigenvector (PC1) from principal component analysis on correlation Hi-C matrix as described previously^85^ with minor modifications. We used HOMER software with parameters ‘-res 500000 -window 1000000’ to obtain the PC1 value based on Pearson Correlation Coefficients (PCC) Hi-C matrix. Sometimes the entry signs of PC1 need to be inverted to ensure that we are assigning the correct signs to individual regions. As GC content is well correlated with A and B compartments^86^, we calculated the GC content of each region and inverted the eigenvector sign if the average GC content of negative-eigenvector entries is higher than that of positive-eigenvector entries. To obtain the heatmap plot of enrichment of A/B interaction, an A/B compartment profile for each chromosome was then separated into 5 bins: (min to 20th percentile), (20th percentile to 40th percentile), etc. For each pair of bins (25 pairs total), the averaged “O/E” values were then calculated for loci belonging to each pair of bins. The compartment strength was then calculated as (AA+BB)/AB^2^ as done previously^87^. For the error bar in evaluating the compartment strength, we obtained 100 5×5 compartment enrichment matrices by bootstrapping. For each pixel of the 5x5 compartment enrichment map, we took all the O/E values that contributed to this pixel and took a random sample with replacement of the same size that the contributing values. We then proceeded with downstream for each of the 100 reshuffled maps.

TADs and loops annotated in CH12.LX were obtained from^59^ and lifted over to the mm10 genome version using the UCSC genome browser liftOver tool. Aggregated plots of TAD enrichment map were obtained by averaging O/E values over annotated TAD positions at 40kb resolution as previously described^87^. For each domain of length *L*, a map for the region ((*Start - L*) to (*End + L*)) was obtained; this produced a contact that is three times bigger than a given domain. This contact map was then rescaled to a (90×90) pixel map using linear interpolation and block-averaging. In the resulting map, the mid-region pixels 30 to 60 correspond to the TAD body. TAD strength for boxplots was quantified as the ratio of two numbers. The first number is the within-TAD intensity: the sum of the central square of the enrichment map, rows 30 to 59 and columns 30 to 59. The second number is the between-TAD intensity, ½ of the sums of the regions [0:30, 30:60] and [30:60, 60:90]. Aggregated plots of loop enrichment map were obtained by averaging O/E values around loop anchors of 310kb window at 10kb resolution. Loop strength was calculated as previous described^87^.

Hi-C “de novo boundary” aggregate plots at MERVLs are centered on 5’ to 3’-oriented MERVL and show a window of 2mb around the MERVL element at 40kb resolution. For illustrating the change of insulation around MERVL genes, the log_2_ transformed Hi-C O/E matrix was further scaled by *z*-score normalization across each row. We calculated the insulation score as originally defined^60^ with minor modifications. Briefly, for each region i in the genome, we calculated the average number of O/E interactions in 40kb Hi-C matrix in a quadratic window with the lower left corner at (i-1, i+1), and the top right corner at (i-5, i+5), where 5 is the window size in bins. We normalized insulation scores by dividing each region’s score by the average scores of the nearest 50 regions, and log_2_-transforming the resulting vector, thus accounting for local biases in insulation score. Visualization of Hi-C matrix was carried out by Juicer tool (https://github.com/aidenlab/juicer)88 and R software (https://www.r-project.org/). For heatmap visualization in Juicer, we converted valid read pairs into .hic format files with the juicer tool pre command.

### Statistical analysis

All statistical analyses for Next Generation Sequencing (NGS) data were performed with R/Bioconductor software utilizing custom R scripts. The other statistical analyses were performed with GraphPad Prism software. Details of individual tests are outlined within each figure legend, including number of replications performed (n) and the reported error as standard error of the mean (s.e.m). All statistics are * p < 0.05, ** p < 0.01, *** p < 0.001, and were calculated by Wilcox signed rank test (for paired samples), Mann-Whitney U test (for independent samples), two-way ANOVA as described in the figure legends.

### Code availability

All the analysis in this study was made based on custom perl and R codes and can be available upon reasonable request.

### Data availability

All the bulk RNA-seq, single-cell RNA-seq, ChIP-seq, ATAC-seq and Hi-C data generated in this study have been deposited in the National Center for Biotechnology Information Gene Expression Omnibus (GEO) database under the accession number of GSE166041. Previously published RNA-Seq data that were re-analyzed here are available under accession codes GSE33923, GSE51682, GSE74278, GSE85632, GSE100939, GSE120953, GSE113671, GSE97778 and GSE66582. Published ChIP-Seq data for DUX are available under accession code GSE85632. Published ChIP-Seq data for p53 are available under accession code GSE26360. Published ATAC-seq data are available under accession code (GSE66390 and GSE85632). Published Hi-C data of mouse pre-implantation embryos are available under accession code GSE82185. Published Hi-C data of lymphoblastoid cells are available under accession code GSE63525. Supplementary Table 5 provided a summary for all analyzed NGS datasets used in this study. All other data supporting the findings of this study are available from the corresponding author on reasonable request.

## Acknowledgements

We thank Dr. Xiong Ji for providing the Pol I degraded mES cell lines. We thank Dr. Pengxu Qian for providing the snoRNA KO cell lines. We thank Dr. Todd Macfarlan for kind discussion. J.Z. is supported by the National Key Research and Development Program of China (No. 2018YFA0107100, No. 2018YFA0107103, No. 2018YFC1005002), the National Natural Science Foundation projects of China (No. 31871453, No. 91857116) and the Zhejiang Natural Science Foundation projects of China (No. LR19C120001). H.Y. is supported by the Zhejiang Natural Science Foundation projects of China (No. LQ21C120002).

## Author contributions

J.Z. and H.Y. conceived and designed the study and experiments. H.Y. and J.Z. wrote the manuscript with contributions from all authors. H.Y. designed and performed all computational analysis. Z.S., T.T., H.P., A.L., Y.Z., L.C. and L.Z. performed the molecular experiments in mES cells. J.Z., L.Z. and J.C. performed the experiments in mouse embryos. H.Y. and Y.X. assisted with the experiment sample preparation. M.C., S.G. and G.D provided computational support and gave critical suggestions about the study design and paper writing. All authors analyzed the results and approved the final version of the manuscript.

## Additional information

### Competing interests

The authors declare no competing interests.

---

**Extended Data Fig.S1:**
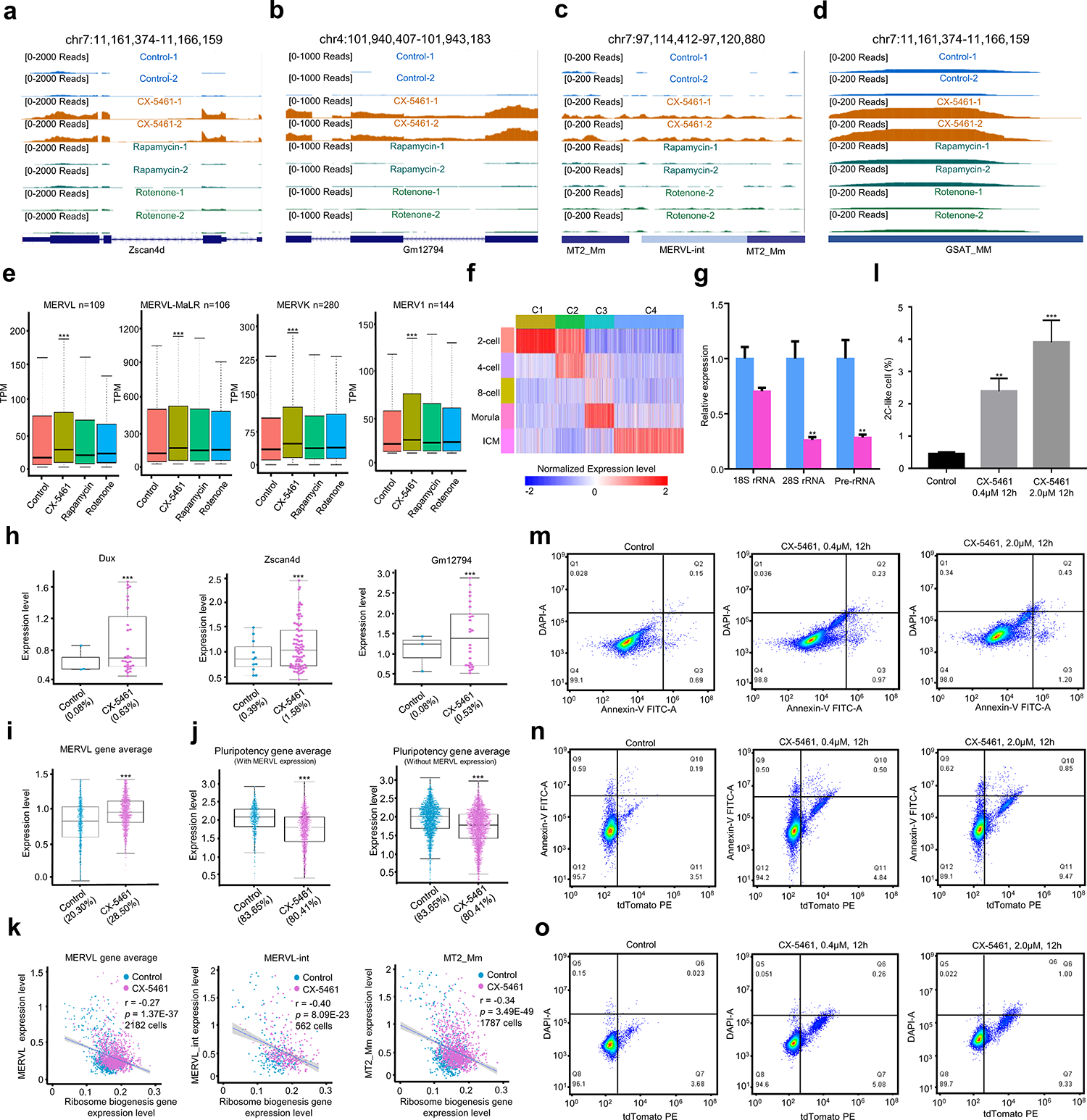
Inhibition of rRNA biogenesis activated 2C-like transcriptional program and induced an expanded 2C-like cell population in mES cells, related to Fig.1. **a)** UCSC Genome Browser viewing of RNA-sequencing results in *Zscan4d* gene locus (GEO accession GSE166041). **b)** UCSC Genome Browser viewing of RNA-sequencing results in *Gm12794* gene locus (GEO accession GSE166041). **c)** UCSC Genome Browser viewing of RNA-sequencing results in a representative ERV (*MERVL-int* and *MT2_Mm*) gene locus (GEO accession GSE166041). **d)** UCSC Genome Browser viewing of RNA-sequencing results in a representative *GSAT_MM* gene loci (GEO accession GSE166041). **e)** Boxplots show the expression levels of major ERV gene classes under different types of cellular stress treatment. n denotes the number of sub-classes of ERV genes, ***: p<0.001, Wilcox signed rank test (GEO accession GSE166041). **f)** A heatmap plot demonstrating four developmental stage-specific gene clusters derived from RNA-seq data of pre-implantation mouse embryos (GEO accession GSE97778). **g)** qRT-PCR quantification of rRNA expression in control mES cells and CX-5461 treated mES cells, **: p<0.01, ***: p<0.001, two-way ANOVA, the replicates of experiment n=3. **h)** Boxplots demonstrating the expression levels of 2C marker genes of Dux, Zscan4d and Gm12794 in control mES cells and CX-5461 treated mES cells; Each point denoted a single cell; The number of parentheses denotes the percentage of cells expressing these genes; ***: p<0.001, Wilcox signed rank test (GEO accession GSE166041). **i)** Boxplots demonstrating the averaged expression levels of *MERVL* genes in control mES cells and CX-5461 treated mES cells with single cell RNA-sequencing analysis; Each point denoted a single cell; The number of parentheses denotes the percentage of cells expressing *MERVL* genes; ***: p<0.001, Wilcox signed rank test (GEO accession GSE166041). **j)** The expression levels of pluripotency genes in control mES cells and CX-5461 treated mES cells; Each point denoted a single cell; The number of parentheses denotes the percentage of cells expressing these genes; ***: p<0.001, Wilcox signed rank test (GEO accession GSE166041). **k)** Scatter plots demonstrating negative correlation of expression level between *MERVL/MERVL-int/MT2_Mm* and ribosome biogenesis genes; Each dot represents a single cell with detectable ERV expression; *r* denotes correlation coefficient; *p*-value was obtained by cor.test function in R software (GEO accession GSE166041). **l)** The percentage of 2C::tdTomato positive cells was quantified using FACS analysis in control mES cells and CX-5461-treated mES cells; Data are means ± SD; SD: Standard Deviation; **: p<0.01, ***: p<0.001, two-way ANOVA, the replicates of experiment n=5. **m)** FACS analysis on Annexin-V FITC, marker for early cell apoptosis, and DAPI, marker for late cell apoptosis upon different treatment doses of CX-5461; **n)** FACS analysis on Annexin-V FITC and 2C::tdTomato positive mES cells upon different treatment doses of CX-5461. **o)** FACS analysis on DAPI and 2C::tdTomato positive mES cells upon different treatment doses of CX-5461. In **e)**, **h)**, **i)** and **j)**, the centre of the box plots represents the median value and the lower and upper lines represent the 5% and 95% quantile, respectively.

**Extended Data Fig.S2:**
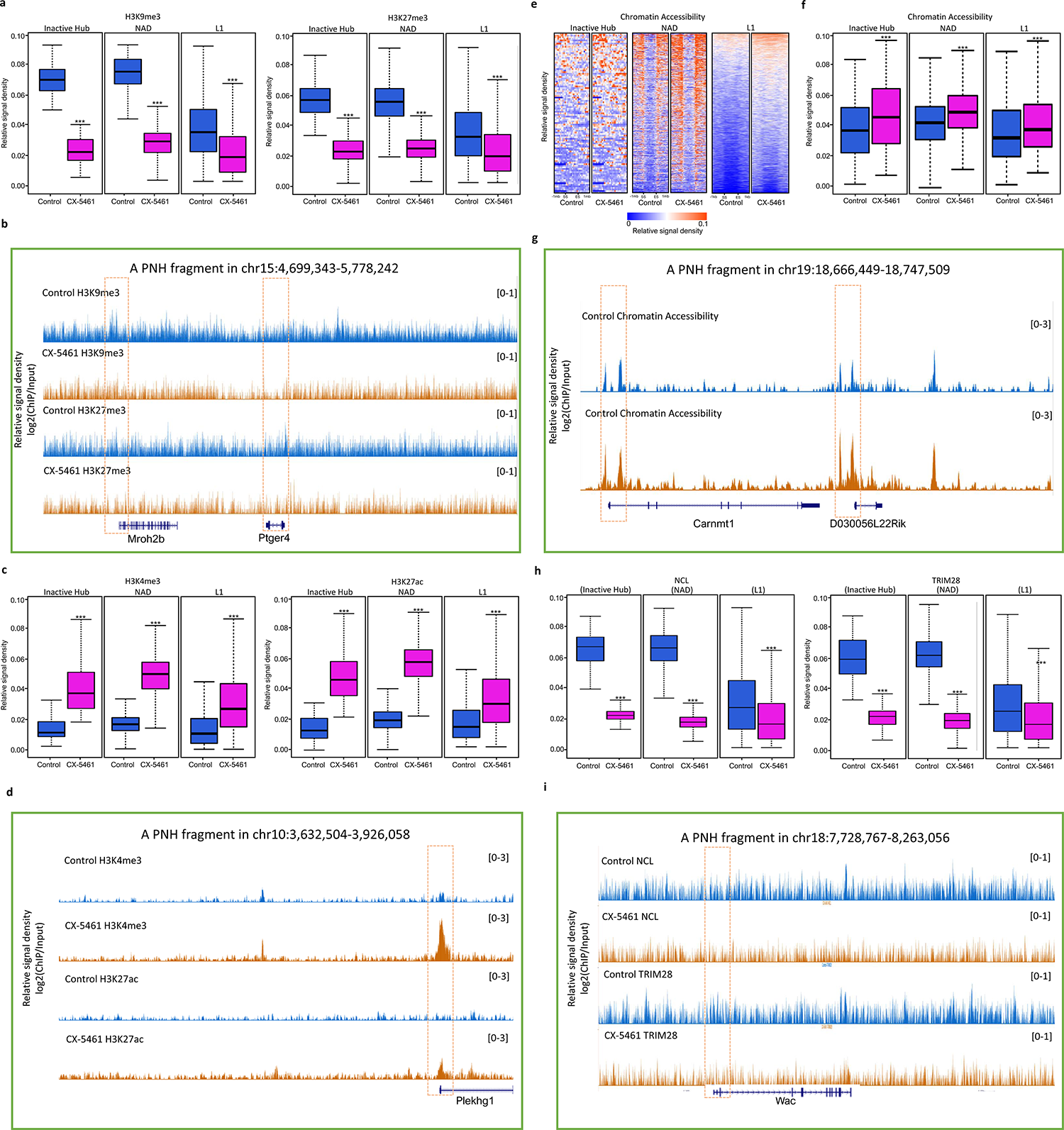
Deficiency of rRNA biogenesis disrupted normal nucleolar LLPS and epigenetic state of PNH region, related to Fig.2. **a)** Boxplots demonstrate the averaged H3K9me3 and H3K27me3 levels of 101 Inactive Hub fragments, 578 NAD fragments and 34888 L1 sequences; ***: p<0.001, Wilcox signed rank test (GEO accession GSE166041). **b)** UCSC Genome Browser viewing of H3K9me3 and H3K27me3 ChIP-seq signals in control and CX-5461 treated mES cells around a representative PNH fragment at chr15:4,699,343-5,778,242 (GEO accession GSE166041). **c)** Boxplots demonstrate the averaged H3K4me3 and H3K27ac levels of 101 Inactive Hub fragments, 578 NAD fragments and 34888 L1 sequences; ***: p<0.001, Wilcox signed rank test (GEO accession GSE166041). **d)** UCSC Genome Browser viewing of H3K4me3 and H3K27ac ChIP-seq signals in control mES cells and CX-5461 treated mES cells around a representative PNH fragment at chr10:3,632,504-3,926,058 (GEO accession GSE166041). **e)** Heatmap plots demonstrate the levels of chromatin accessibility on within 1mb region around start and end sites of Inactive Hub and NAD, and within 1kb region around start and end sites of L1 (GEO accession GSE166041). The regions of different lengths of Inactive Hub and NAD fragments were fitted to 1mb. The regions of different lengths of L1 sequences were fitted to 1kb. SS: start site of a chromatin fragment of PNH; ES: end site of a chromatin fragment of PNH. The PNH fragment was defined as the L1 contained regions overlapped with Inactive Hub and NAD. **f)** Boxplots demonstrate the averaged chromatin accessibility of 101 Inactive Hub fragments, 578 NAD fragments and 34888 L1 sequences; ***: p<0.001, Wilcox signed rank test (GEO accession GSE166041). **g)** UCSC Genome Browser viewing of ATAC-seq signals in control and CX-5461 treated mES cells around a representative PNH fragment at chr19:18,666,449-18,747,509 (GEO accession GSE166041). **h)** Boxplots demonstrate the averaged binding signals of NCL and TRIM28 on 101 Inactive Hub fragments, 578 NAD fragments and 34888 L1 sequences; **: p<0.01, ***: p<0.001, Wilcox signed rank test (GEO accession GSE166041). **i)** UCSC Genome Browser viewing of NCL and TRIM28 ChIP-seq signals in control and CX-5461 treated mES cells around a representative PNH fragment at chr18:7,718,063-8,074,255 (GEO accession GSE166041). In **a)**, **c)**, **f)** and **h)**, the centre of the box plots represents the median value and the lower and upper lines represent the 5% and 95% quantile, respectively.

**Extended Data Fig.S3:**
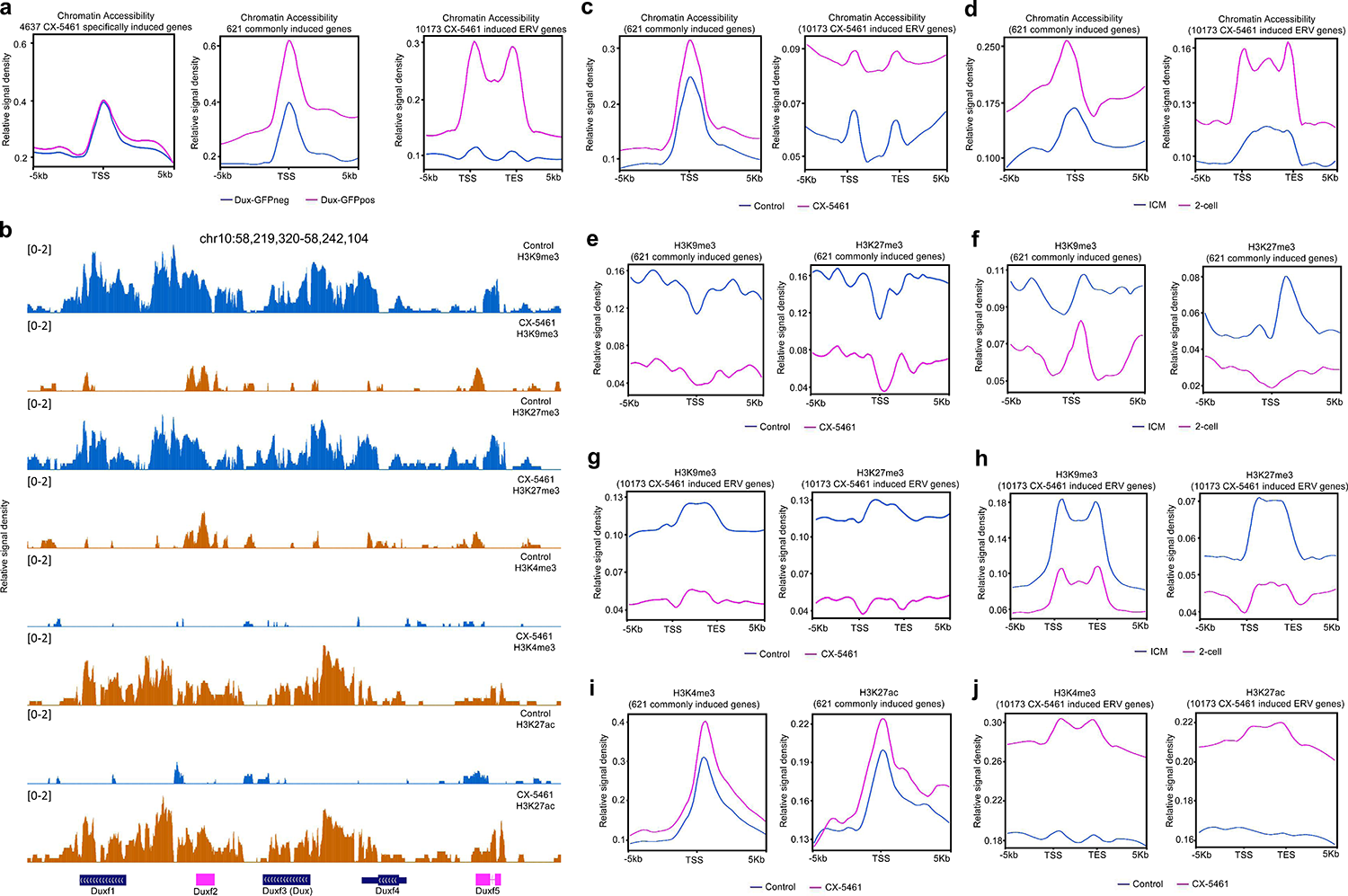
2C/ERV genes were activated through Dux, related to Fig.3. **a)** Line plots demonstrate the meta-analysis results of chromatin accessibility in Dux-GFP positive mES cells and Dux-GFP negative mES cells within 5kb region around transcription start sites or transcription start and end sites of 621 commonly induced genes between CX-5461 treatment and *Dux* overexpression, 4637 specifically induced genes by CX-5461 and 10173 CX-5461 induced ERV genes using published ATAC-seq data. The regions of different lengths of ERV genes were fitted to 5kb (GEO accession GSE85632). **b)** UCSC Genome Browser viewing of ChIP-seq signals around *Dux* locus (GEO accession GSE166041). **c)** Line plots demonstrate the meta-analysis results of chromatin accessibility of 621 commonly induced genes between CX-5461 treatment and *Dux* overexpression and 10173 CX-5461 induced ERV genes in control mES cells and CX-5461 treated mES cells. The regions of different lengths of ERV genes were fitted to 5kb (GEO accession GSE166041). **d)** Line plots demonstrate the meta-analysis results of chromatin accessibility of 621 commonly induced genes between CX-5461 treatment and *Dux* overexpression and 10173 CX-5461 induced ERV genes in 2-cell embryo and ICM embryo. The regions of different lengths of ERV genes were fitted to 5kb (GEO accession GSE66390). **e)** Line plots demonstrates the meta-analysis results of H3K9me3 and H3K27me3 levels of 621 commonly induced genes between CX-5461 treatment and *Dux* overexpression in control mES cells and CX-5461 treated mES cells (GEO accession GSE166041). **f)** Line plots demonstrate the meta-analysis results of H3K9me3 and H3K27me3 levels of 621 commonly induced genes between CX-5461 treatment and *Dux* overexpression in 2-cell embryo and ICM embryo (GEO accession GSE66390). **g)** Line plots demonstrate the meta-analysis results of H3K9me3 and H3K27me3 levels of 10173 CX-5461 induced ERV genes in control mES cells and CX-5461 treated mES cells; The regions of different lengths of ERV genes were fitted to 5kb (GEO accession GSE166041). **h)** Line plots demonstrate the meta-analysis results of H3K9me3 and H3K27me3 levels of 10173 CX-5461 induced ERV genes in 2-cell embryo and ICM embryo; The regions of different lengths of ERV genes were fitted to 5kb (GEO accession GSE66390). **i)** Line plots demonstrate the meta-analysis results of H3K4me3 and H3K27ac levels of 621 commonly induced genes between CX-5461 treatment and *Dux* overexpression in control mES cells and CX-5461 treated mES cells (GEO accession GSE166041). **j)** Line plots demonstrate the meta-analysis results of H3K4me3 and H3K27ac levels of 10173 CX-5461 induced ERV genes in control mES cells and CX-5461 treated mES cells; The regions of different lengths of ERV genes were fitted to 5kb (GEO accession GSE66390).

**Extended Data Fig.S4:**
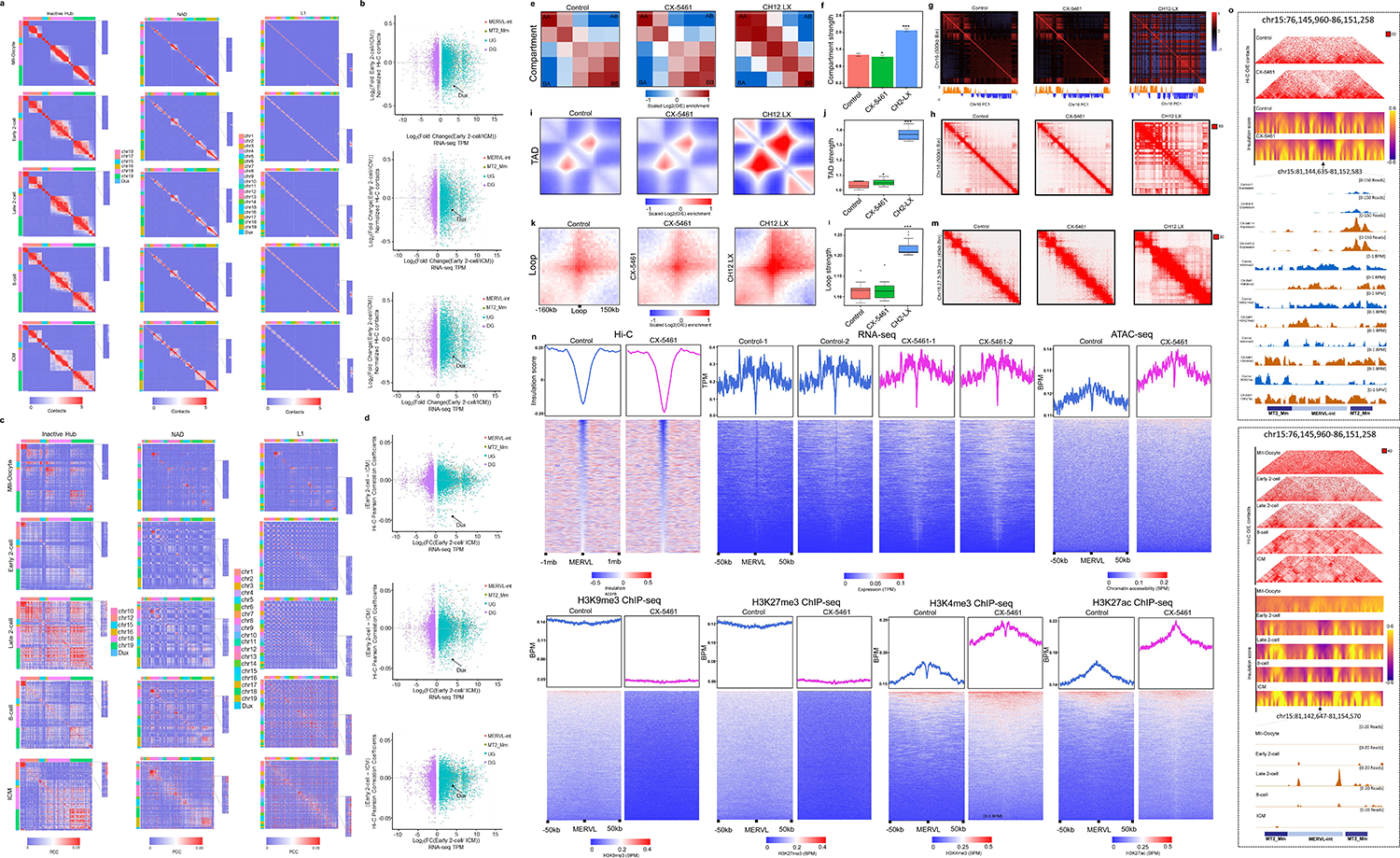
rRNA biogenesis defect drove 3D chromatin structure reorganization of PNH and MERVL regions towards the 2C-like state, related to Fig.4. **a)** Hi-C contact maps of Inactive Hub/NAD/L1 and 1.5 Mb genomic regions around *Dux* at 150kb resolution during mouse pre-implantation embryos development. The zoomed-in regions aim to demonstrate the change of Hi-C contacts between *Dux* and chromosome 10 during mouse pre-implantation embryos development (GEO accession GSE82185). **b)** Scatter plot demonstrates the log_2_(fold change) of Hi-C contacts between Inactive Hub/NAD/L1 and different types of genes in early 2-cell and ICM stage embryos (GEO accession GSE82185). **c)** Hi-C pearson correlation heat maps of Inactive Hub/NAD/L1 and 1.5 Mb genomic regions around *Dux* at 150kb resolution during mouse pre-implantation embryos development. The zoomed-in regions aim to demonstrate the change of Hi-C PCC between *Dux* and chromosome 10 during mouse pre-implantation embryos development (GEO accession GSE82185). **d)** Scatter plot demonstrates the PCC difference between Inactive Hub/NAD/L1 and different types of genes in early 2-cell and ICM stage embryos (GEO accession GSE82185). **e)** A/B interaction profile showing contact enrichment between active and inactive compartments (GEO accession GSE166041 and GSE63525). **f)** Quantification of compartment strength; *: p<0.05, ***: p<0.001, Wilcox signed rank test. **g)** Pearson correlation heat maps for chromosome 16 at 500kb resolution to demonstrate A/B compartment (GEO accession GSE166041 and GSE63525). **h)** Hi-C contact maps for chromosome 16 at 500kb resolution for A/B compartment profile. **i)** Observed/Expected (O/E) aggregate plot of TADs (GEO accession GSE166041 and GSE63525). **j)** Quantification of TAD strength; *: p<0.05, ***: p<0.001, Wilcox signed rank test (GEO accession GSE166041 and GSE63525). **k)** O/E aggregate plots of chromatin loops (GEO accession GSE166041 and GSE63525). **l)** Quantification of loop strength; ***: p<0.001, Wilcox signed rank test (GEO accession GSE166041 and GSE63525). **m)** Hi-C contact maps for chromosome 16:27.3-36.2mb region at 40kb resolution to demonstrate TAD and chromatin loop (GEO accession GSE166041 and GSE63525). **n)** Insulation score, expression (TPM), ATAC-seq (BPM), H3K9me3 (BPM), H3K27me3 (BPM), H3K4me3 (BPM) and H3K27ac (BPM) signals of control and CX-5461-treated mES cells centered on CX-5461-induced *MERVL* genes (GEO accession GSE166041). **o)** Representative 40kb Hi-C O/E interaction matrices of a *MERVL* loci (chr15:76,145,960-86,151,258) located at TAD boundaries are shown as heatmaps, along with insulation score and genome browser tracks of RNA-Seq, H3K9me3, H3K27me3, H3K4me3 and H3K27ac ChIP-Seq signals of the expanded genomic region containing the TAD boundary (arrows) in control and CX-5461-treated mES cells as well as in mouse early embryos (GEO accession GSE166041).

**Extended Data Fig.S5:**
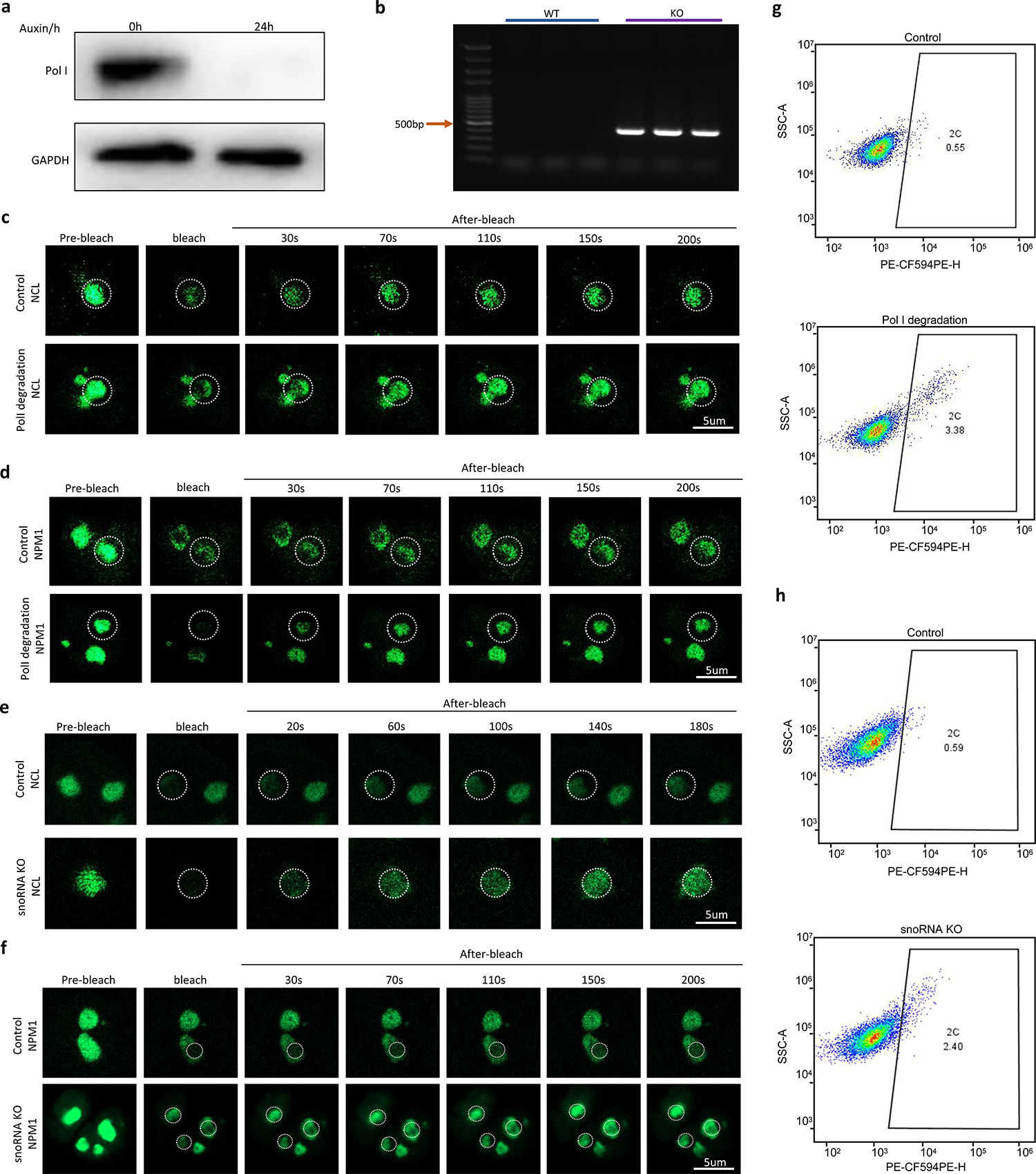
Genetic interferences of rRNA biogenesis recapitulate CX-5461-induced 2C-like molecular phenotypes, related to Fig.5. **a)** Western Blotting experiment showing the Pol I protein degradation after 24h of Auxin treatment. **b)** PCR experiment showing that a 400bp band was observed in the snoRNA KO mES cells, but not in the wild-type (WT) mES cells. As a band of 400bp was designed especially in the snoRNA KO mES cells, this result indicates that the homologs of human SNORD113-114 gene cluster was successfully knocked-out. **c)** Shown images are representative of 4 times of NCL FRAP experiments in control mES cells and Pol I degraded mES cells. **d)** Shown images are representative of 4 times of of NPM1 FRAP experiments in control mES cells and Pol I degraded mES cells. **e)** Shown images are representative of 4 times of NCL FRAP experiments in control mES cells and snoRNA knockout mES cells. **f)** Shown images are representative of 4 times of NPM1 FRAP experiments in control mES cells and snoRNA knockout mES cells. **g)** FACS analysis on 2C::tdTomato+ mES cells in Pol I degraded mES cell lines, showing the change of percentage of 2C-like cells. **h)** FACS analysis on 2C::tdTomato+ mES cells in snoRNA knockout mES cell lines, showing the change of percentage of 2C-like cells.

**Extended Data Fig.S6:**
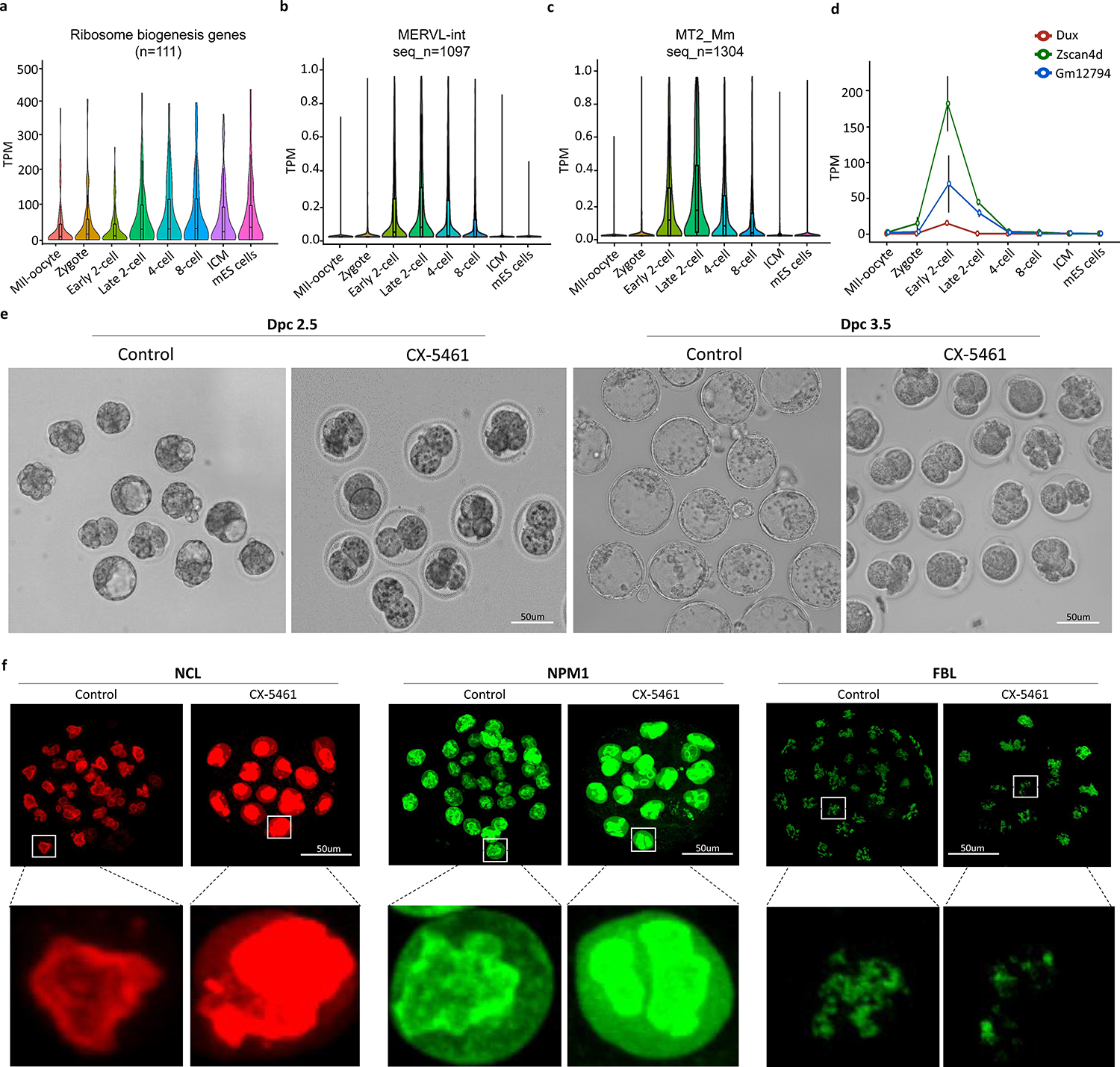
rRNA biogenesis is critically required at the 2-cell-to-4-cell stage transition during pre-implantation embryo development, related to Fig.6. **a)** Expression pattern of Ribosome biogenesis gene set across different early embryo developmental stages. n denotes the number of sub-classes of *MERVL* genes. **b)** Expression pattern of *MERVL-int* genes across different embryo developmental stages. seq_n denotes the number of annotated *MERVL-int* sequences in the mouse mm10 reference genome. **c)** Expression pattern of *MT2_Mm* genes across different embryo developmental stages. seq_n denotes the number of annotated *MT2_Mm* sequences in the mouse mm10 reference genome. **d)** Expression pattern of 2C marker genes, *Dux*, *Zscan4d* and *Gm12794*, across different embryo developmental stages. **e)** Representative images of mouse embryos produced from control and CX-5461 treatment during two different developmental stages; Dpc: Days post-coitum. This experiment was repeated three times independently with similar results. **f)** Immunofluorescence staining of NCL, NPM1 and FBL in control blastocyst embryos and CX-5461-treated blastocyst embryos.

## Notes

### Competing Interest Statement

The authors have declared no competing interest.

